# Protein-Like Polymer for Inhibition of Tau Fibril Propagation in Human-Derived Models of Neurodegeneration

**DOI:** 10.1101/2025.02.02.636155

**Authors:** Andrew P. Longhini, Mara Fattah, Julia Sala-Jarque, Austin DuBose, Erica Keane Rivera, Olivia R Sclafani, Jessica Zhou, Jacqueline E. Anatot, Matthew P. Thompson, Matthew T. Unger, Emily R. Beckett, Colby Fagan, Dylan Nakamura, Richard I. Morimoto, Songi Han, Baofu Qiao, Nathan C. Gianneschi, Kenneth S. Kosik

**Author notes:** Corresponding authors (A.P.L.), (N.C.G.), (K.S.K.). These authors contributed equally to this work (Andrew P. Longhini, Mara Fattah).

## Abstract

The misfolding, aggregation, and spread of tau protein fibrils underlie tauopathies, a diverse class of neurodegenerative diseases for which effective treatments remain elusive. Among these are corticobasal dementia (CBD) and progressive supranuclear palsy (PSP), canonical examples of 4-repeat (4R) tauopathies characterized by tau isoforms exclusively with four microtubule-binding repeat domains. We target this 4R tau isoform-specific mechanism by focusing on misfolded tau’s distinctive stem-loop-stem structural motif formed by the junction of the 4R-defining alternatively spliced exon and the adjacent constitutive exon. A synthetic peptide based on this stem-loop-stem sequence can induce aggregation and spread in an isoform-specific manner. Here, we develop a protein-like polymer (PLP) in which multiple copies of this synthetic peptide form a brush-like structure capable of preventing tau aggregation by binding and capping fibril ends *in vitro*, in human brain organoids, and in cellular models with an EC50 of 105 ± 14 nM. PLPs demonstrate robust activity against fibrils derived from CBD and PSP patient brains and a PS19 mouse tauopathy model. Previous tau-targeted treatments have primarily focused on broad tau clearance, aggregation inhibition, or microtubule stabilization, often lacking isoform specificity and precision. In contrast, this approach targets the 4R tau isoform’s unique structural motif, offering a tailored therapeutic intervention for diseases like CBD and PSP. Supported by prior studies showing blood-brain barrier penetrance and safety profiles, this tau-binding PLP offers a promising translational path toward clinical applications in tauopathy treatment.

## Introduction

Protein misfolding and aggregation are hallmarks of numerous neurodegenerative and systemic diseases, in which aggregates spread through a prion-like mechanism, templating healthy proteins to adopt pathological conformations ^1–6^. Diseases such as Alzheimer’s ^7,8^, Parkinson’s ^3^, amyotrophic lateral sclerosis (ALS)^9^, and systemic amyloidosis^6^ exhibit this pattern of progressive protein aggregation as fibrillar seeds grow, adding proteins to their propagating face. As these misfolded, aggregated proteins accumulate, they impose an increasing burden on cellular degradation pathways, such as the ubiquitin-proteasome system and autophagy, which may already be compromised due to aging ^10–12^. The inability of these systems to clear aggregates allows their unchecked growth, resulting in cellular dysfunction, toxicity, and ultimately cell death ^12–14^. A therapeutic approach aimed at targeting the propagating face of these fibrils, thus blocking the further recruitment and conversion of unaffected protein, holds promise in halting the spread of aggregates by preventing their elongation, thereby mitigating disease progression.

Tauopathies represent specific neurodegenerative disorders characterized by the prion-like spread of pathological tau aggregates through interconnected neuronal networks^1,4,15^. Each tauopathy has characteristic structural folds, as determined by cryo-TEM single particle reconstruction, and distinct anatomical distributions ^16–23^. Given tau’s central role in disease progression, targeting misfolded tau spread and seeding is a promising strategy for disrupting disease progression. We recently described a 19-residue peptide containing a P301L mutation and spanning the R2/R3 splice junction of tau. This peptide folds and stacks into seeding-competent fibrils capable of specifically inducing aggregation of 4R, but not the 3R isoform of tau. These tau peptide fibrils propagate aggregated intracellular tau over multiple generations of cell division ^24^. The peptide adopts a U-shaped fold, closely aligning with the structures that define glial tauopathy-progressive supranuclear palsy-tau (GPT), corticobasal degeneration (CBD), and progressive supranuclear palsy (PSP) 4R tauopathies ^25^. Notably, the peptide acts as a mini-prion, accurately mimicking 4R isoform-specific spread of tau in cellular models.

Small peptides have been identified which are capable of acting as capping agents, preventing the elongation and propagation of tau aggregates ^26–34^. During disease progression, tau fibrils facilitate misfolded transitions by adding naïve tau monomers. These fibrils can then fragment, exit the originating cells, and spread to neighboring cells, thereby advancing disease ^35,36^. To prevent this templating and spread, efforts have focused on designing peptides that effectively bind to and “cap” the growing fibril ends, hindering further aggregation ^26–34^. While this technique has shown promise, it remains in its nascent stages.

A significant challenge with peptides is their rapid degradation and limited cellular penetration, not to mention that they generally lack the ability to traverse the blood-brain barrier ^37^. Concurrently, there has been focus on developing antibodies and nanobodies that inhibit tau spread. Such designs incorporate capping peptides directly into the variable CD3 domain of nanobodies ^38^. Indeed, our own previous work demonstrated that the VHH-Z70 nanobody significantly reduced the seeding of naïve tau in culture systems ^24^.

To overcome these obstacles, we utilized a synthetic proteomimetic strategy that incorporates peptide side chains as high-density brush polymers termed protein-like polymers (PLPs, **Figure 1)** ^39–44^. PLPs incorporating hydrophobic polymer backbones adopt a globular conformation, which imbues the side chains with resistance to proteolytic degradation while retaining and enhancing the bioactivity of the conjugated peptides via multivalent display. This multivalency leads to PLPs typically exhibiting on the order of 100-1000 times the binding constant of peptides alone ^39,42^. The inherent features of transient amphiphilicity (metaphilicity) driving cell penetration, facile peptide modification, and multivalency combine to make PLPs suitable for cell entry, cytosolic distribution, and engagement of intracellular target proteins like tau ^45–47^.

In this paper, we introduce a novel approach to target tau spread using protein-like polymers (PLPs), functionalized with peptides designed to target tau fibrils. These functionalized PLPs halt the growth of tau fibrils *in vitro* and effectively prevent the seeding of naïve tau by preformed tau fibrils from both disease and synthetic sources in cells. Our findings have substantial implications for tau-based therapeutics, as these functionalized PLPs are not only multi-functional but have also been shown to exhibit favorable pharmacokinetic properties ^47^.

### Results

We have previously identified a 19-amino acid peptide that could recapitulate 4R tauopathy disease folds, as determined by cryo-TEM single particle reconstruction (**Figure 1A**) ^24^. These results suggested an opportunity for tau isoform-specific therapeutic interventions. Published work on the jR2R3 P301L fibril, which mimics 4R conformation-specific tauopathies such as CBD and PSP, led us to postulate that a jR2R3 P301L peptide — containing the aggregation-prone PHF6 motif — could bind and cap a propagating fibril if displayed as a PLP. We modified this peptide to prepare functional PLPs and named it the tau-binding peptide (TBP). We hypothesize that while a monomeric peptide of jR2R3 P301L can readily add to a growing end of a preformed fibril seed, when presented on a globular, multivalent PLP further tau monomer addition would be occluded as the symmetry of the propagating face would be broken, giving rise to an effective capping agent (**Figure 1B**).

### Synthesis and characterization of Tau PLPs and control PLPs

The peptide sequence for the TBP-PLP was modified based on design considerations to permit cellular uptake, improve water solubility, and space the binding domain away from the backbone allowing key residues to interact with tau. Therefore, the original jR2R3 P301L sequence (DNIKHVLGGGSVQIVYKPV) was augmented with cationic charge residues (**Figure 1A**). Specifically, C-terminal lysines were incorporated into TBP-PLP to provide hydrophilicity and cell penetration potential ^40^. While the peptide monomer already incorporates a flexible hydrocarbon linker to connect the norbornene polymerizable unit to the peptide, additional PGGSK residues were appended to the DNIK sequence of the jR2R3 P301L variant (**Figure 1A**). The modification was designed to increase conformational flexibility and ensure optimal accessibility of the binding region to tau protein targets, and mimics part of the native tau sequence (**Figure 1B**).

PLP syntheses were performed as previously described (see Supporting Information) ^45–47^. Briefly, peptides were synthesized via solid-phase peptide synthesis wherein the addition of a norbornene polymerizable unit (N-(hexanoic acid)-exo-5-norbornene-2,3-dicarboximide) was conjugated as a non-natural amino acid to the N terminus of the peptide (**Figure 1C**). Peptide monomers were cleaved from resin as described in Methods, purified via preparative high-performance liquid chromatography (prep-HPLC), and further characterized by electrospray ionization mass spectrometry (ESI-MS) and analytical HPLC for molecular weight determination and purity (**Figures S1**) ^48,49^.

Peptide monomers were polymerized using ring-opening metathesis polymerization (ROMP), and the kinetics of polymerization were monitored for completion via nuclear magnetic resonance spectrometry (NMR, **Figure S2**). Upon consumption of the peptide monomer, a norbornene-conjugated cyanine 5.5 (Cy5.5) dye label was incorporated into the polymer backbone before termination if a dye labeled version of the PLP was desired (TBP-PLP-Cy5.5, **Figure S3**). For some studies, TBP-PLP was additionally biotinylated with a biotin-terminating agent previously reported (TBP-PLP-biotin, **Figures S4, S5**) ^50^.

Polymers lacking labels were terminated with ethyl vinyl ether (EVE) and purified via prep-HPLC. The resulting pure fractions were lyophilized to obtain the PLPs in powder form. PLPs were characterized to determine molecular weight using sodium dodecyl-sulfate polyacrylamide gel electrophoresis (SDS-PAGE) and size exclusion chromatography multi-angle light scattering (SEC-MALS) (**Figures S3**, **S4**). These PLPs were designed to contain 15 repeats of the modified TBP peptide (**Figure 1D**). An additional two PLPs, TBP-Keap1-PLP and TBP-Keap1-PLP-Cy5.5, were prepared as tau binders and Keap1 recruiters for targeted tau degradation via Keap1-driven ubiquitination.

**Figure 1:**
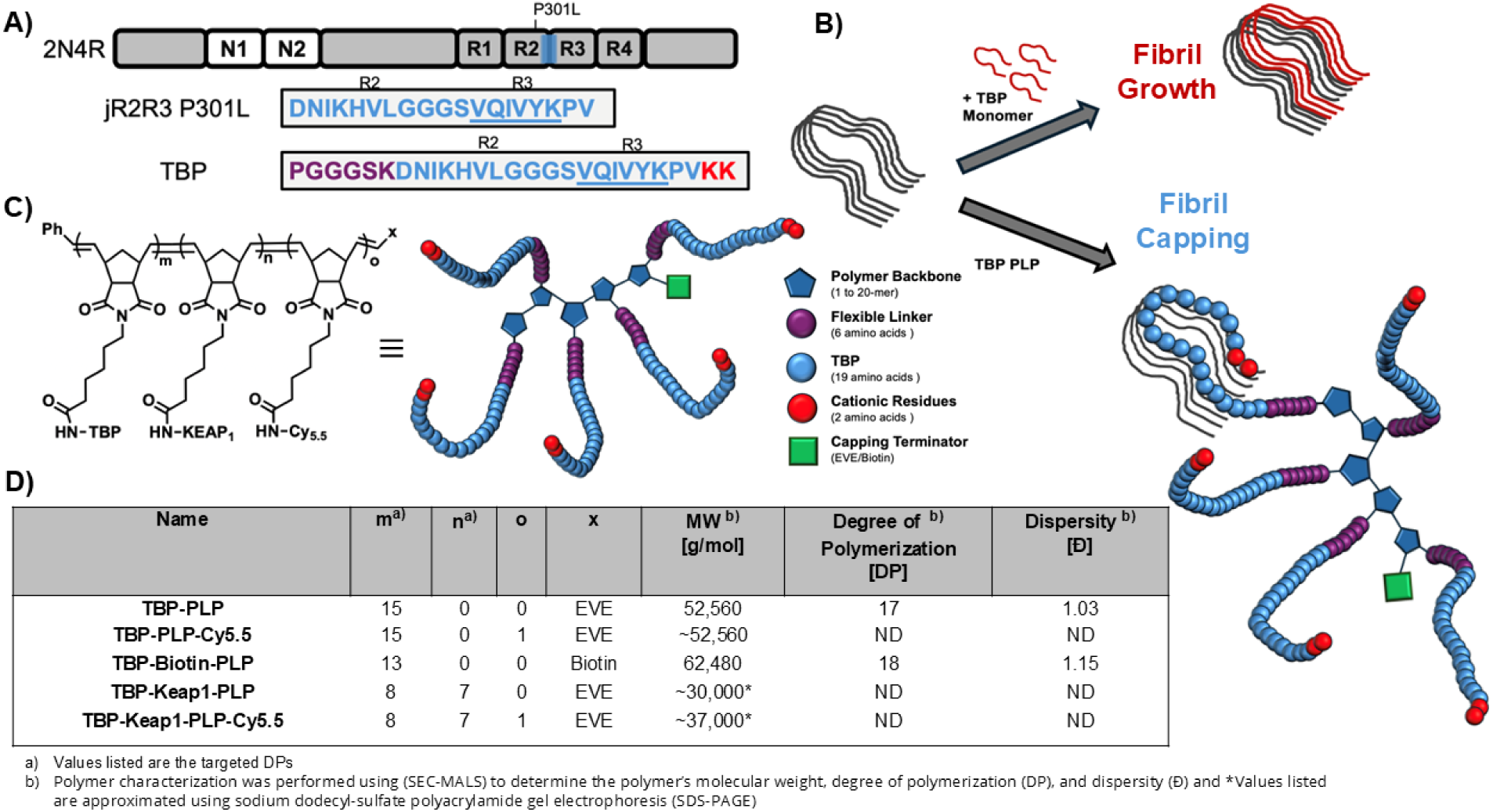
Tau Binding Peptide Fragment Selection for Polymerization, Polymer Design, and Fibril Capping Mechanism. A) Overview of tau-originating peptide (jR2R3 P301L, blue) modified to form a new tau-binding peptide (TBP) with the PHF6 sequence underlined. To increase its flexibility at the PLP backbone, a linker sequence was added (purple) and to increase its solubility and cell penetrance two C-terminal lysines were added (red) B) Tau fibrils can seed the fibrillization of naïve tau monomers (top) or TBP-PLP binding can cap and inhibit fibril growth. C) General structure of the PLPs, with simplified scheme showing the flexible linker in purple, the jR2R3 P301L sequence in blue, and the C-terminus lysines in red. An optional terminating group containing a fluorophore or biotin is shown as a green box. D) Table showing PLPs used in this study. EVE = ethyl vinyl ether.

### TBP-PLP prevents tau seeding *in vitro*

TBP-PLP was tested for efficacy in inhibiting tau fibril formation *in vitro*. Our previous work demonstrated that jR2R3 P301L fibrils can seed the formation of new fibrils from jR2R3 P301L monomers when fibrils are introduced at low concentrations ^24^. To quantify this in the presence of TBP-PLP, 25µM jR2R3 P301L monomer was added to 25µM jR2R3 P301L fibrils (with fibril concentration based on the concentration of individual peptides in the fibril solution). Fibril formation was then monitored via transmission electron microscopy (TEM) and Thioflavin T (ThT) fluorescence assays, evaluating the process across TBP-PLP concentrations ranging from 3.125 to 25µM (**Figures 2A, 2B**).

In the absence of TBP-PLP, fibril formation occurred rapidly, reaching a gradually increasing plateau by 4 hours, as indicated by a sharp rise in ThT fluorescence. In contrast, TBP-PLP was introduced at concentrations as low as 6.25µM, there was a near-complete suppression of ThT fluorescence, indicating strong inhibition of fibril formation. All concentrations above 6.25µM led to complete inhibition of fibril formation as monitored by both TEM and ThT fluorescence (**Figures 2A, 2B**).

TEM images comparing reactions at 0µM and 6.25µM TBP-PLP (Figure 2A) showed the formation of long paired-helical fibrils in the absence of the PLP. In contrast, samples treated with TBP-PLP displayed a notable absence of fibrils; the few fibrils that were present appeared clumped at the thin filamentous ends (**Figure 2A, Bottom**). These results suggest that TBP-PLP prevents fibril elongation through a steric hindrance mechanism as hypothesized, wherein the PLP binds to the fibril ends, blocking further addition of monomeric tau.

**Figure 2:**
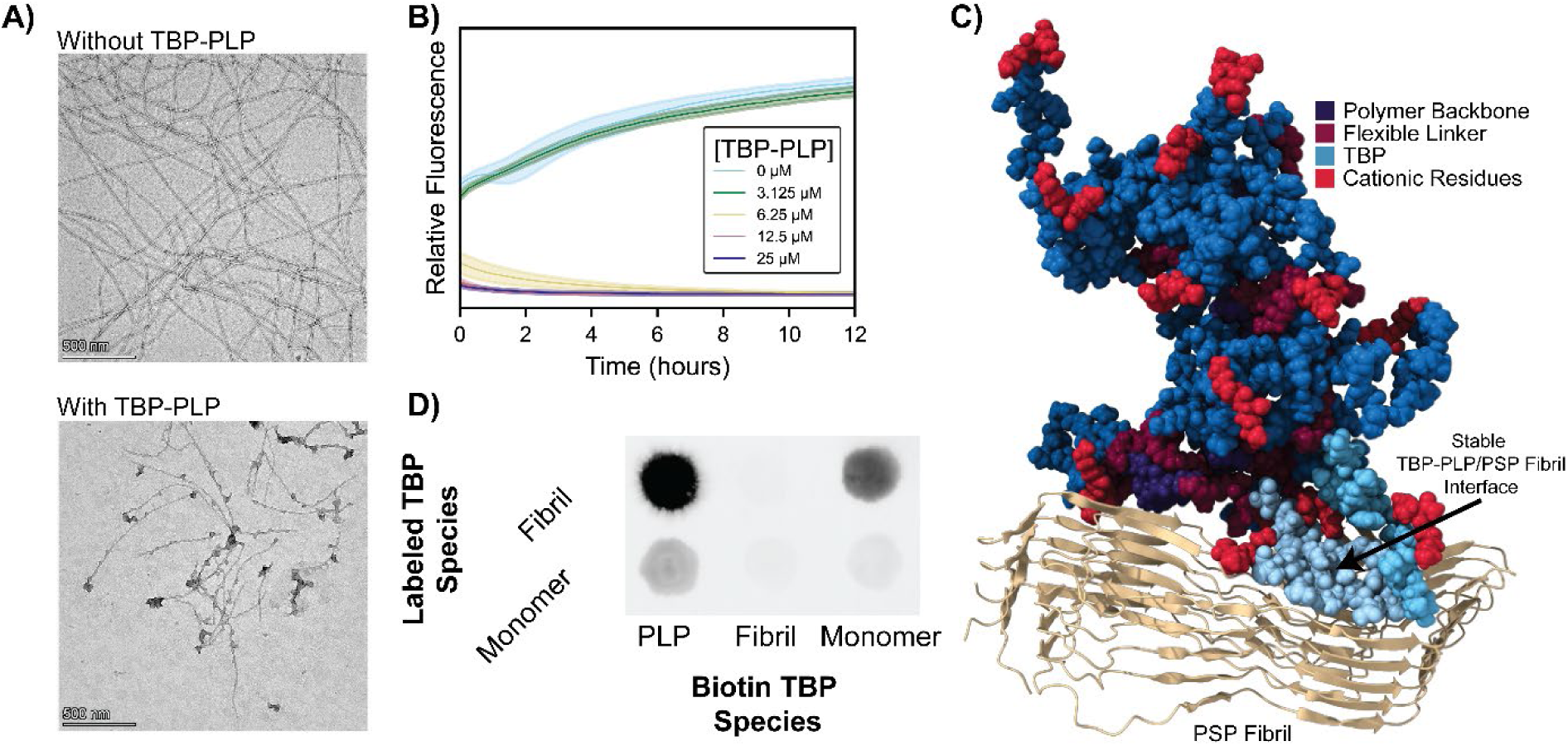
TBP-PLP Prevents jR2R3 P301L Fibril Formation. A) jR2R3 P301L fibrils were used to seed monomeric jR2R3 P301L peptide. In the absence of TBP-PLP, the formation of long paired-helical filaments was observed as previously reported ^24^. In the presence of TBP-PLP, a marked absence of filaments was observed, and those that were present exhibited large globular structures decorating their ends. Scale Bar: 500nm. B) Increasing concentrations led to a complete absence of fibril formation as reported by a ThT assay. C) Simulations of TBP-PLP and PSP fibril (PDB:7P65) ^21^ show a stable interface between the fibril and the PLP arms that caps and prevents further fibril elongation. D) TBP-PLP-biotin, jR2R3 P301L-biotin fibrils, or jR2R3 P301L-biotin monomers were incubated with fluorescently tagged jR2R3 P301L-Cy5 fibril or jR2R3 P301L-Cy5 monomer. Pull-downs of the biotin-tagged species with streptavidin beads showed that the TBP-PLP-biotin interacted with both Cy5-labeled fibrils and monomers, jR2R3 P301L-biotin fibrils pulled down small quantities of Cy5 labeled monomer or other Cy5-labeled fibrils, and jR2R3 P301L-biotin monomers pulled down Cy5-labeled fibrils and to a much lesser extent other Cy5-labeled monomers.

### Molecular Dynamics Reveal TBP-PLP-Capping Mechanism Inhibiting Tau Fibril Elongation

Motivated by our hypothesis that TBP-PLP can occlude the growing face of tau fibrils, we performed coarse-grained (CG) molecular dynamics simulations using the polarizable MARTINI 2.2P force field^51,52^ and GROMACS ^53^. By employing CG simulations, we were able to sample microsecond timescales (20 μs) on a system comprising the tau fibril structure (PDB: 7P65) ^21^ and the TBP-PLP in an aqueous environment with 0.14 M NaCl. This setup allowed us to overcome the size and complexity constraints typical of all-atom simulations.

During the CG simulation, the TBP-PLP monomers were initially free in solution near the fibril. Over the course of 20 μs, a single TBP-PLP monomer spontaneously adsorbed onto the fibril’s amyloidogenic jR2R3 region, forming stable β-sheet interactions (**Figure S6**, **7**). Notably, these interactions closely matched the complementary binding interface at the fibril’s “growing face,” suggesting TBP-PLP could act as a physical cap that prevents naïve tau monomers from docking. These findings established a mechanistic rationale for how TBP-PLP might sterically block fibril extension, consistent with our *in vitro* TEM observations.

To further characterize the stability and molecular details of this CG-identified contact, we next performed all-atom explicit-solvent MD simulations using the CHARMM 36m potential ^54^ in GROMACS. Given the large conformational space and extended timescales needed for a spontaneous approach, we initiated the simulation with one TBP-PLP peptide side chain positioned at the CG-derived fibril-binding region (**Figure S8A**). After a short equilibration phase with minimal position restraints (to preserve the initial alignment), we released these restraints and ran an unrestrained simulation for 300ns (**Figure S8B**).

Strikingly, the initial TBP-PLP:fibril β-sheet interaction remained stable throughout the entire simulation, reinforcing the CG prediction that the fibril face is an energetically favorable binding site for the TBP-PLP. Moreover, an additional peptide chain from the same TBP-PLP molecule began to stack atop the first, further occluding the fibril-binding interface. Snapshots from the trajectory reveal that these TBP-PLP segments extend outward from the fibril surface, creating a steric barrier that would likely hinder naive full-length tau monomers from initiating the critical “touchdown” step required for fibril elongation (**Figure 2C**).

Together, the coarse-grained and all-atom simulations suggest a model in which TBP-PLP monomer arms bind and cap the growing fibril face, effectively blocking the initial docking of naive tau monomers. By forming stable β-sheet interactions with the amyloidogenic region and stacking additional peptide segments over the interface, the TBP-PLP substantially reduces solvent accessibility and sterically hinders further fibril extension. This mechanism aligns with the reduced fibril growth observed in our in vitro assays and provides a structural basis for the observed inhibition of tau aggregation.

### TBP-PLP Interacts Strongly with jR2R3 P301L Monomers and Fibrils

To further characterize the interaction between TBP-PLP and tau, we used a combination of dot-blot assays and isothermal titration calorimetry (ITC) to directly measure the binding of TBP-PLP to monomeric and fibrillar tau species.

We first biotin-tagged the jR2R3 P301L monomer, fibril, and TBP-PLP constructs. In parallel, we prepared fluorescently labeled stocks of the jR2R3 P301L monomer and fibril, maintaining a 1:10 ratio of labeled to unlabeled species. Equimolar concentrations (10µM) of the biotin-labeled species and fluorescently labeled species were incubated together for 1 hour at 37°C. The mixtures were then incubated with streptavidin-coated magnetic beads for an additional 30 minutes at 37°C. Following stringent washing, bound species were eluted and detected via dot-blot assays based on their fluorescent signals.

The results showed that TBP-PLP-biotin bound robustly to both fluorescently labeled monomers and fibrils, with a stronger signal observed in the case of fibrils. This enhanced signal is in part explained by the fact that a single fibril can contain multiple fluorophores, whereas a monomer can only contain one. Consequently, the binding of a single fibril leads to a greater overall fluorescence signal. In contrast, biotin-tagged fibrils showed minimal interaction with other fluorescently labeled fibrils or monomers. However, biotin-tagged monomers pulled down fluorescently labeled fibrils, but showed much weaker interaction with monomers (**Figure 2D**). Control experiments, where fluorescently labeled species were incubated with streptavidin beads in the absence of a biotin-labeled species, produced no detectable signal in the elution (**Figure S9**).

To quantify the interaction of TBP-PLP with monomers, we employed isothermal titration calorimetry (ITC). In these experiments, 500 nM TBP-PLP was titrated with 5µM jR2R3 P301L monomer. The interaction generated an exothermic reaction, and fitting the data to an independent binding model yielded a dissociation constant (K_D_) of 18 nM, indicating strong binding (**Figure S10**). However, attempts to derive a binding constant for jR2R3 P301L fibrils were unsuccessful due to the complexity of the interaction. Although we know the monomer concentration of the fibrils and that each fibril has two ends, the length of the fibrils is highly variable. Further, because we cannot determine how many monomers each fibril contains, it is not possible to accurately calculate the fibril end concentration, making it challenging to derive a K_D_ for this interaction.

### TBP-PLP Directly Enters Cells in Culture Models

To characterize TBP-PLP cell uptake, we measured the fluorescence intensity over time following PLP administration, normalizing to total cellular area to account for differences in cell density and size across trials (**Figure 3**). Concentrations ranging from 0.25 to 2 μM were tested, and results indicated that the cellular uptake of PLP increased with concentration until a plateau was observed at approximately six hours post-administration at all concentration ranges (**Figure 3A**). This plateau suggests a saturation point in the uptake mechanism, which is consistent with direct cellular penetration rather than receptor-mediated endocytosis, as the latter would likely display a more gradual, sigmoidal uptake pattern due to the involvement of cellular machinery like receptors and vesicle formation. This is consistent with behavior seen for other PLPs, where uptake is proportional to concentration within the same time frame^45,46^, and when a singular concentration produces the same uptake signals between 2 and 24 hours, suggesting saturation.

Additionally, a linear correlation between PLP concentration and uptake at the 6-hour timepoint (R²=0.95, Figure 3A, inset) further supports the hypothesis of passive entry. Endocytosis, which is an active and regulated process that typically involves vesicle formation and trafficking, does not scale linearly with increasing ligand concentration. This deviation from expected endocytic kinetics reinforces the idea that PLPs penetrate the cell membrane directly, bypassing these more regulated pathways. Imaging cells treated with fluorescently labeled PLP at six hours, showed a population of cells in which a clear PLP signal is associated with the cell membrane before becoming a cytoplasmic signal (**Figure 3B**).

To further assess whether endocytosis plays a role in PLP uptake, we next examined the colocalization of the labeled PLPs with markers for key endocytic compartments: Rab5 for endosomes and Lamp1 for lysosomes (**Figures 3C**). Cells were treated with labeled PLPs for six hours, fixed, and stained with antibodies specific to these endosomal markers. No significant colocalization was observed between the PLPs and any of these markers, suggesting that the PLPs do not enter the canonical endocytic pathway (**Figure 3C**). The lack of colocalization with Rab5 also indicates that PLPs do not undergo vesicular trafficking via endosomes, further supporting the hypothesis of a non-endocytic mode of entry.

To confirm the independence of PLP uptake from traditional endocytic pathways, we treated the cells with inhibitors of key components of endocytosis. Cytochalasin D, which disrupts actin polymerization, was used to inhibit clathrin-mediated endocytosis, while filipin III, which sequesters cholesterol, was employed to block caveolae-mediated uptake. Dynasore, a dynamin inhibitor, was used to prevent scission of endocytic vesicles. Despite these treatments, no reduction in PLP uptake was observed after four hours (**Figure 3D**). The persistence of robust PLP internalization under these conditions reinforces the conclusion that these PLPs penetrate cells through a mechanism independent of the endocytic machinery. We hypothesize this is due to the metaphilic behavior of PLPs ^55,56^—a phenomenon describing the transient amphiphilicity of the PLP, driven by the combination of a lipophilic polymer backbone and hydrophilic terminal cationic residues, which enable engagement with and passage through the lipid membrane^45^.

These results collectively support the conclusion that TBP-PLP directly penetrates cell membranes in a concentration-dependent manner, bypassing the endocytic pathway. This cell-penetrating ability opens up many therapeutic applications by enabling cytosolic delivery and circumventing the limitations of endocytic trafficking, which often leads to degradation and entrapment in lysosomes.

**Figure 3:**
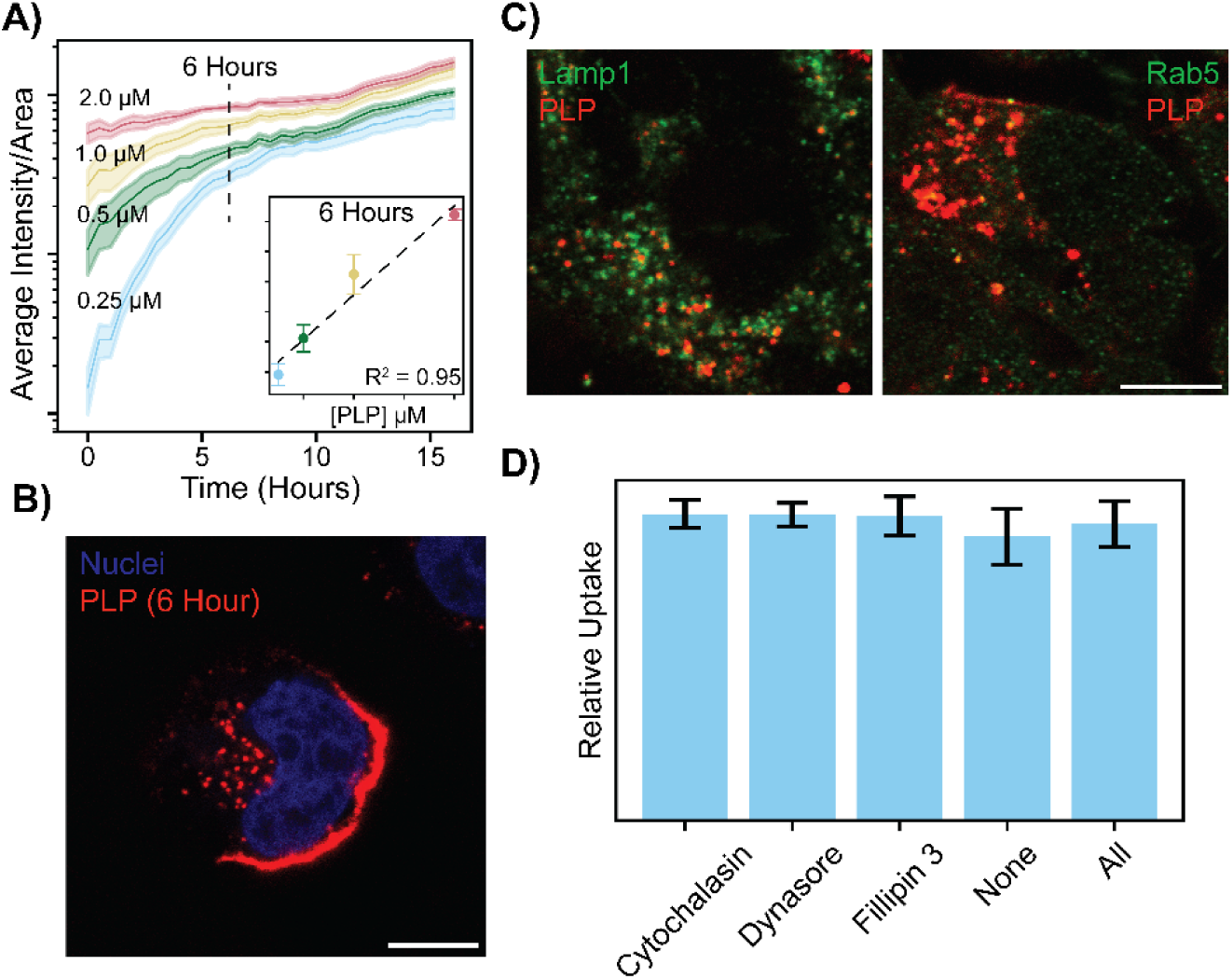
Uptake of TBP-PLP Through Direct Cellular Penetration. A) Increasing concentrations of TBP-PLP-Cy5.5 (0.25µM—cyan, 0.5µM—green, 1µM—sand, 2.0µM—rose) were monitored by live cell imaging every 30 minutes over 16 hours. The average intracellular fluorescence intensity normalized by total cellular area was plotted with standard error of the mean (SEM) against time (hours). A dotted line at 6 hours indicates the approximate point where the log plot plateaus, signifying a change in uptake rate. The inset shows a strong linear relationship (R²=0.95) between peptide concentration and uptake at the zero time point. B) Representative live cell image at 6 hours post-PLP exposure, showing intracellular PLP accumulation and membrane-associated PLP (red). Scale bar: 10µM. C) Co-staining of PLP (red) with Lamp1 (green, left) and Rab7 (green, right) showing no colocalization between PLP and these endocytic markers. Scale bar: 10µM. D) Cells were treated with cytochalasin D, dynasore, filipin III, or all inhibitors in combination, and PLP uptake was imaged at 24 hours. PLP uptake remained unaffected across all treatments, confirming the endocytosis-independent mechanism of cellular entry.

### TBP-PLPs Prevent Seeding of jR2R3 P301L Fibrils in a Cellular Culture System

We previously constructed a cell line expressing a fluorophore-tagged tau construct, mClover3-Tau187 P301L, to monitor the seeding activity of exogenous tau fibrils. Using this model, jR2R3 P301L fibrils strongly induced tau aggregation in a 4R isoform-specific manner ^24^. Extending our *in vitro* ThT findings, we hypothesized that TBP-PLP could effectively inhibit the seeding of jR2R3 P301L fibrils in our cell culture models. In this assay, TBP-PLP was added for six hours, then washed out of the media before the addition of jR2R3 P301L fibrils (**Figure 4A**). We reasoned that removing the TBP-PLP before fibril addition would allow only the internalized PLP to interact with the fibrils, thereby preventing seeding while still facilitating cellular uptake.

We began our investigation by adding 1 μM FITC-labeled jR2R3 P301L fibrils and 2 μM TBP-PLP-Cy5.5 to wild-type cells that do not express mClover3-Tau187-P301L, and monitored their colocalization (**Figure 4B**). Robust colocalization of fibrils and PLP was observed, with nearly all fibrils colocalized with PLP (Manders’ M1 = 0.99 ± 0.004). This high M1 value indicates a strong overlap of PLP with the fibrils. However, there was a significant amount of PLP that did not colocalize with the fibrils, as suggested by the M2 value (Manders’ M2 = 0.46 ± 0.10), indicating that approximately 54% of the PLP signal was not associated with the fibrils. Sixteen biological replicates were included in the colocalization analysis.

For inhibition studies, we treated cells with TBP-PLP at concentrations ranging from 62.5 nM to 4µM for six hours, followed by a media change to prevent extracellular interactions with the subsequently added fibrils. Imaging conducted 24 hours after fibril addition revealed a potent, sub-µM inhibitory effect in the presence of TBP-PLP, with an EC50 of 105 ± 14 nM (R² = 0.91) (Figure 4C, S12A). In contrast, a control protein-like polymer incorporating a previously designed capping peptide ^26^, VQPINK-PLP, failed to block seeding activity, underscoring the effectiveness of our rationally designed construct against 4R mimetic mini-prion fibrils (**Figure S11**).

Further, we introduced the TBP alone, not conjugated to the PLP, into cell cultures (**Figure S11C, D).** These peptides, containing the same sequence used on the PLP conjugates, did not prevent tau aggregation, highlighting that the three-dimensional, multivalent display exhibited by the jR2R3 P301L motif on the PLP scaffold is crucial for effectively blocking further aggregate growth in our cellular model.

To assess long-term inhibition, we imaged cells 48 hours post-fibril exposure. After an initial six-hour incubation with TBP-PLP, cells were rinsed and exposed to fibrils. At 48 hours, aggregate formation increased compared to the 24-hour mark, with the percentage of cells containing aggregates rising from 1.2 ± 0.8% at 4µM PLP to 22.9 ± 7.3% (**Figure 4C, S11B**). Although there was a decrease in the fit to a dose-response model (R² = 0.67), the EC50 was still 479 ± 100 nM compared to 105 ± 14 nM at 24 hours. We hypothesized that at these extended time points, either the peptides conjugated to the PLP are being degraded, or the introduced fibrils are being degraded, leading to the release of newly available fibril ends that are free to seed naïve tau aggregates.

**Figure 4:**
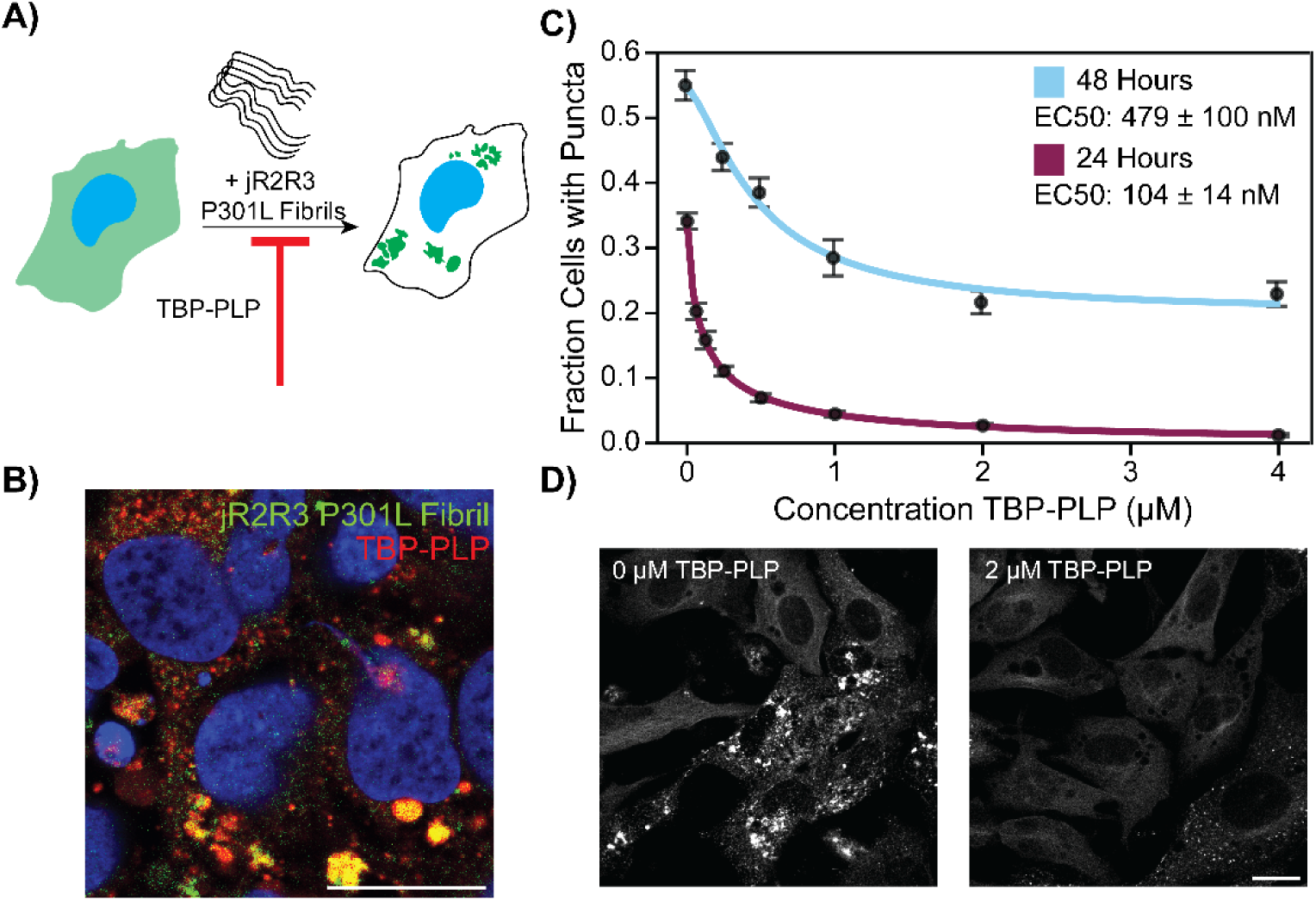
Inhibition of Cellular Tau Aggregates by TBP-PLP. A) Schematic representation of the assay setup. Cells stably expressing mClover3-tau187-P301L were incubated with jR2R3 P301L fibrils in the presence or absence of TBP-PLP. In the absence of PLP, cells accumulated punctate tau inclusions, indicating aggregation. B) TBP-PLP-Cy5.5 (red) and fluorescently labeled jR2R3 P301L fibrils (green) were added to cells and imaged after 24 hours. Scale bar: 20 μM. C) The fraction of cells containing puncta was plotted against the concentration of TBP-PLP (n = 15 per condition). Each point represents the average of independent measurements, with error bars showing the standard error of the mean. The data were fit to dose-response curves. After 24 hours of fibril exposure, the EC50 (the concentration of PLP at which 50% of cells exhibited a reduction in puncta) was calculated to be 104 ± 14 nM, with the percentage of cells displaying puncta approaching 0% at 4 μM TBP-PLP. At 48 hours post-fibril addition, the reduction in puncta was less pronounced, with 20% of cells still displaying puncta and an EC50 of 479 ± 100 nM for the reduction in puncta. D) Representative images of cells treated with 0 μM and 2 μM TBP-PLP. In the 0 μM condition, punctate inclusions are visible, while in the 2 μM TBP-PLP-treated cells, no puncta are observed. Scale bar: 10 μM.

### TBP-PLPs are Stable in Serum and Cell Lysate

To test the stability of TBP-PLP and to rule out the possibility that TBP-PLP moieties were being degraded at the 48-hour time point, we subjected them to a range of *in vitro* conditions designed to mimic physiological environments and test their stability (**Figure S12)**. These data show that the PLP retains its function and integrity over extended periods in biologically relevant environments (sera and cell lysate, 37°C for 7 days) and implies that capping of fibrils by TBP-PLP is thermodynamically stable. Other studies on PLPs stability support these findings of extended stability in solution and under relevant biological conditions ^40,46,47^.

### Prolonged Exposure and Multiple Doses of TBP-PLP Prevent 48-hour Aggregation

Based on the stability of TBP-PLP to a variety of conditions, we believe that the reduced efficacy in preventing seeding at 48 hours is due to the fragmentation of tau fibrils leading to an excess of fibril ends to cap. We speculated that maintaining a constant intracellular concentration of PLP could enhance long-term aggregation prevention. To test this, we implemented two dosing strategies that differed from our initial washout protocol.

In the first approach, TBP-PLP was introduced to the cells for six hours before fibril addition, as in our previous experiments. However, unlike earlier procedures, the PLP was not removed from the media prior to adding the fibrils. This strategy was based on our observation that, although a significant amount of PLP uptake occurred within the first six hours—as demonstrated by the inflection point in Figure 3A—accumulation continued beyond this point. Employing this method, we noted a substantial reduction in aggregate formation at 48 hours, achieving a percentage of cells with aggregates of 15.4 ± 2.1% at 4 μM PLP, with a EC50 of 628 ± 113 nM (R^2^ = 0.79), compared to 19.5 ± 6.2% and 479 ± 100 nM from previous experiments (**Figure 5A, S13A, B**).

Building on this approach, we added a second dose of TBP-PLP at 24 hours. The results were markedly improved, with an EC50 of 381 ± 45 nM (R^2^ = 0.67) and cells exhibiting puncta decreased to ∼4% at 4 μM PLP **(Figure 5A**). These findings indicate that enhancing intracellular retention and concentrations of PLPs could be an effective strategy for long-term prevention of tau aggregation and spread. This dual-dosing regimen maintains TBP-PLP concentration and caps newly fragmented tau fibrils, thereby preventing seeding of naïve monomeric tau.

**Figure 5:**
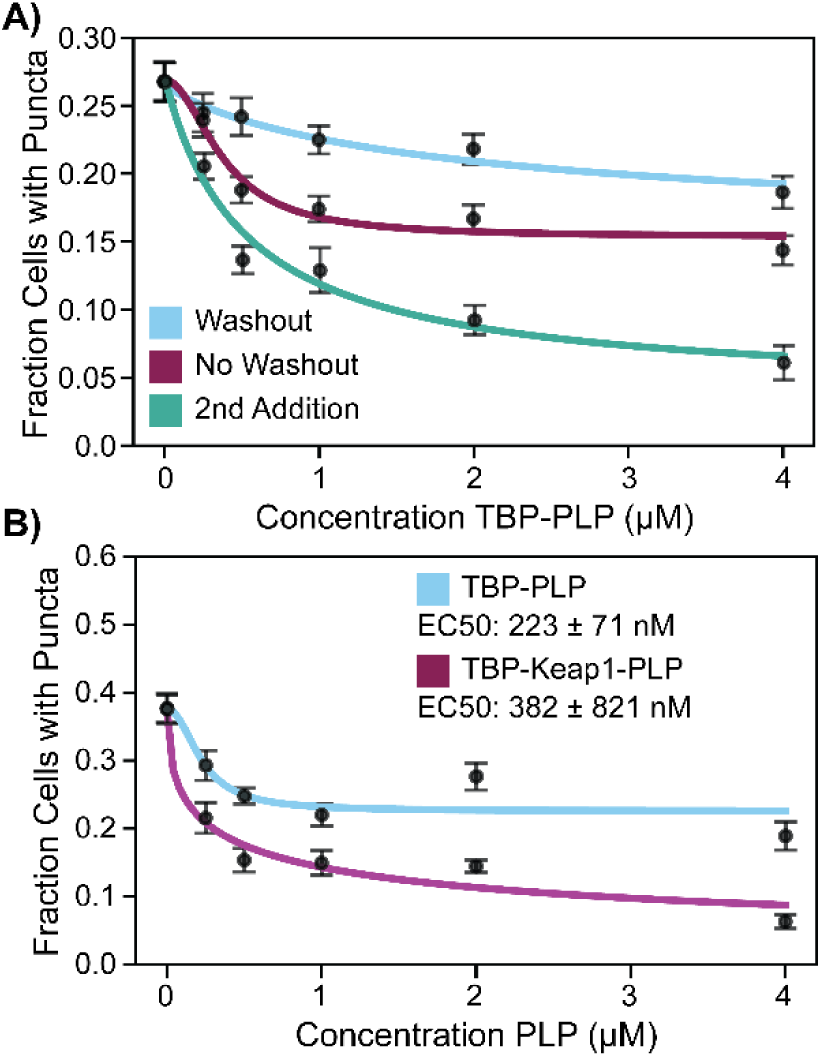
Strategies to Increase the Efficacy of TBP-PLP Over Extended Time Periods. A) TBP-PLP was added to cells under three different conditions: (1) a single 6-hour exposure followed by washout (cyan), (2) a continuous exposure with no washout (purple), or (3) a second dose added at 24 hours (green). Cells were imaged for puncta at 48 hours. Cells receiving two doses (green) exhibited significantly fewer puncta at 48 hours, with only 7.4% ± 2.1% of cells showing fibril formation at a 4 μM concentration, compared to 15% for cells continuously exposed without washout (purple) and 19.5% ± 6.2% for cells exposed for 6 hours followed by wash out (cyan). B) TBP-Keap1-PLP (cyan) demonstrated improved efficacy over the original TBP-PLP (purple) when puncta were analyzed at 48 hours. Next-generation TBP-PLPs that include a Keap1 binding peptide resulted in a more significant reduction in tau aggregates, with cells treated with the Keap1 variant showing a lower percentage of cells with puncta compared to those treated with the TBP-PLP lacking Keap1 recruitment domains.

### Bifunctional PLP for Increased Clearance of Exogenous Tau Fibrils

Since multiple additions of TBP-PLP dramatically increased the effectiveness of aggregate prevention at 48 hours, we hypothesized that directing the PLP-bound fibrils to appropriate degradation machinery could further enhance its ability to prevent tau aggregation. To test this, we engineered a bifunctional PLP containing eight copies of jR2R3 P301L and seven copies of a Keap1-binding peptide, termed TBP-Keap1-PLP. Keap1, when engaged by a binding partner, facilitates the ubiquitination of the binder, thus targeting it for proteasomal degradation.

Previous studies have incorporated a Keap1 peptide sequence successfully into a tau-targeting PROteolysis TArgeting Chimeras (PROTACs)^57^. Keap1 (Kelch-like ECH-associated protein 1) acts as an adaptor protein that facilitates the recruitment of its substrate, Nrf2 (nuclear factor erythroid 2–related factor 2), to a Cullin-RING E3 ubiquitin ligase complex (CRL) that marks the protein for ubiquitination. Given the high affinity of the jR2R3 P301L sequence for tau, we employed it as the foundation for designing a PLP-based PROTAC. Notably, Keap1-binding PLPs have also been explored for engagement and Nrf2 activation in cellular models exhibiting high affinities and potency ^45^. The optimized Keap1-binding sequence, LDPETGEFLRRRR, which carries a net charge of +1, was used in conjunction with the jR2R3 P301L sequence to construct a heterobifunctional PLP PROTAC. This Keap1-binding sequence mimics the binding site of its endogenous partner, Nrf2, enabling selective engagement with Keap1.

In our experiment, both TBP-Keap1-PLP and TBP-PLP were washed out after six hours to assess their lasting effects on tau aggregate prevention. Despite having fewer tau binding sites than the original TBP-PLP, the TBP-Keap1-PLP reduced the percentage of cells with puncta to 5.9% ± 6.0%, compared to 19.5 ± 6.2% for TBP-PLP (**Figure 5B, S13C**). These results suggest that the addition of the Keap1-binding domain effectively enhances the clearance of tau aggregates by directing them to the proteasomal degradation pathway.

Taken together, these findings demonstrate that multifunctional PLPs, such as the PROTAC-like TBP-Keap1-PLP, represent a promising strategy for improving the efficacy of aggregate prevention, providing a dual mechanism of action by both inhibiting aggregation and promoting the degradation of tau fibrils.

### Prevention of Seeding by 4R Tauopathy Patient-Derived and Mouse-Derived Fibrils

The jR2R3 P301L peptide was designed to structurally mimic the fibrils associated with 4R tauopathies. Comparative analysis of the cryo-TEM-derived structures from patient-derived CBD and PSP, as well as PS19 mouse models, with our own jR2R3 P301L fibrils revealed a common strand-loop-strand architecture (**Figure 6A**). This motif was absent in mixed tauopathies such as Alzheimer’s disease (AD), suggesting a structural divergence (Figure 7A). Further, replica exchange molecular dynamics (REMD) simulations of the jR2R3 P301L monomer showed a sampling of various conformations that we predict are capable of binding to disease fibrils (**Figure 6B**). Based on these observations, we hypothesized that TBP-PLP would effectively inhibit seeding by CBD, PSP, and PS19 fibrils, but show reduced efficacy against AD fibrils.

**Figure 6:**
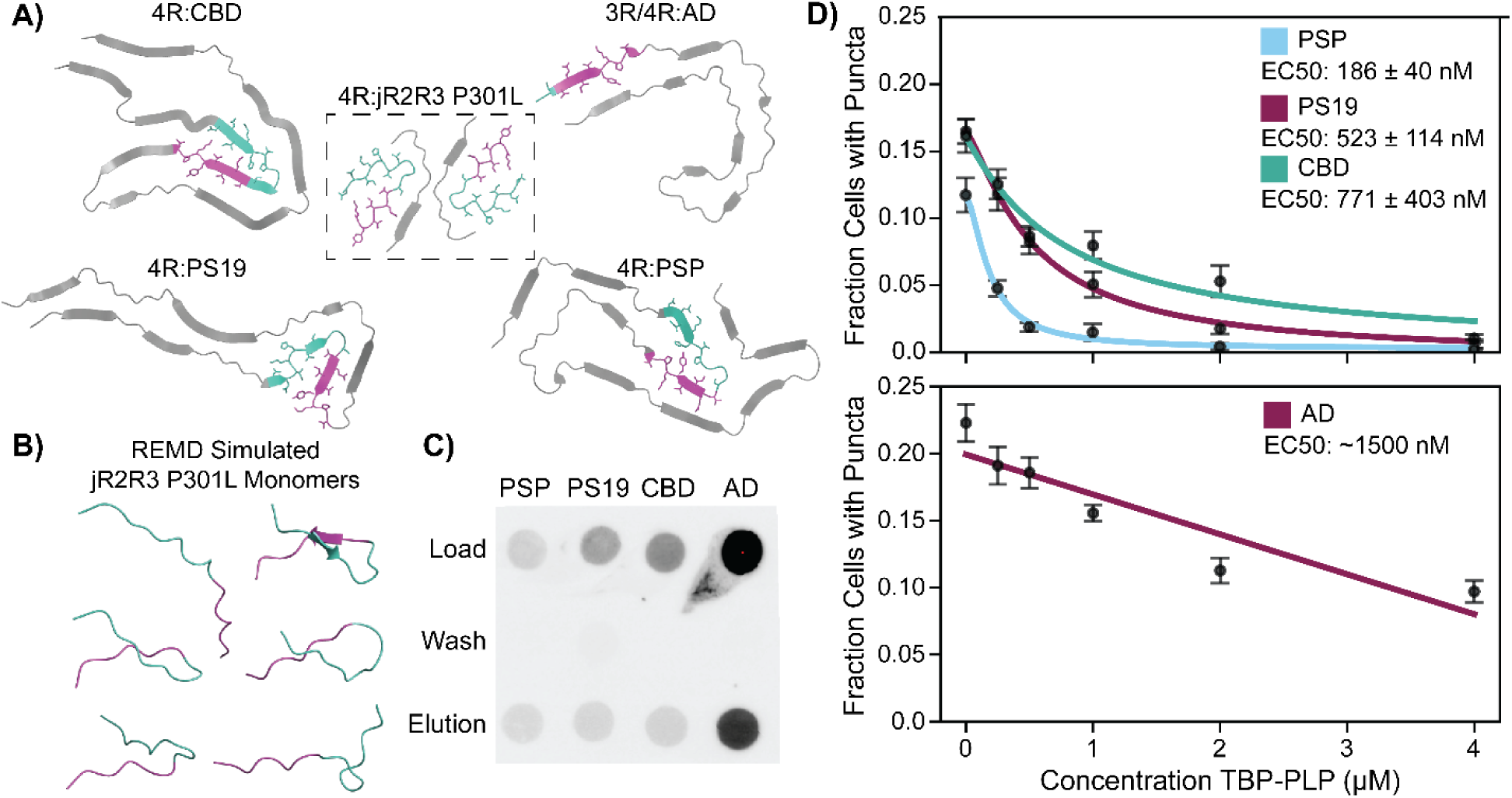
TBP-PLP Prevents Patient-Derived Fibril Seeding. A) Comparison of tau fibril structures from various diseases to the cryo-EM structure of jR2R3 P301L fibrils. The characteristic stem-loop-stem motif observed in 4-repeat (4R) tauopathies, such as CBD (PDB: 6VHA), PS19 (PDB: 8Q92), and PSP (PDB: 7P65), is spanned by the jR2R3 P301L fragment (R2: green, R3: purple). This motif is absent in Alzheimer’s disease (AD) fibrils (PDB: 6VHL). B) Replica exchange simulations of jR2R3 monomers show that the free peptide explores various conformations that can interact with 4R fibrils. C) Dot-blot analysis using TBP-PLP-biotin incubated with magnetic streptavidin beads bound to either PSP, PS19, CBD, or AD fibrils. The complexes were washed, eluted, and detected using a total tau antibody (Tau-5). D) Fibrils from 4R tauopathies (PSP, cyan; PS19, purple; CBD, cyan) were added to cells treated with increasing concentrations of TBP-PLP. The fraction of cells with puncta was plotted against the concentration of TBP-PLP, revealing a characteristic dose-response curve (Upper). In contrast, mixed 3-repeat (3R)/4-repeat (4R) tau AD fibrils (purple) were added to cells treated with increasing concentrations of TBP-PLP. While the fraction of cells with puncta decreased with higher doses, it did not approach baseline levels, and the decrease was unexpectedly linear (Lower). All data points represent at least 15 independent measurements from a minimum of 100 cells each.

To test this hypothesis, we extracted sarkosyl-insoluble material from the brains of patients with CBD, PSP, and AD—as well as material from 9-month-old PS19 mouse brain—and were able to observe fibrils by TEM (**Figure S14**). Dot-blot assays using TBP-PLP-biotin pulled down with magnetic streptavidin beads confirmed specific interactions with the fibrils from CBD, PSP, PS19, and, unexpectedly, with AD fibrils (**Figure 6C, S15**). Fibrils bound selectively to TBP-PLP, with no interaction observed between the fibrils and the streptavidin beads alone. This selective binding supports our hypothesis and underscores the potential of TBP-PLP as a targeted therapeutic approach in distinct tauopathies.

In our cell-based aggregation assay, we optimized experimental conditions to observe a significant proportion of cells with tau aggregation after fibril addition, with PS19 and AD showing aggregation after 24 hours and CBD and PSP requiring 48 hours. These values were determined empirically and will vary based on different fibril preparations from different patients. Our experimental protocol involved treating cells with PLP for six hours before fibril addition without a subsequent washout, to maintain consistent conditions between the 24- and 48-hour time points.

For each of the 4R tauopathies (CBD, PSP, and PS19), we observed a robust prevention of aggregation, characterized by a classic dose-response curve (**Figure 6D, Top, Figure S13D-F**). The EC_50_s for CBD, PSP, and PS19 were 771 ± 403 nM (R^2^ = 0.61), 186 ± 40 nM (R^2^ = 0.73), and 523 ± 114 nM (R^2^ = 0.75), respectively. In contrast, the dose-response curve for AD did not exhibit a typical logarithmic trend but was instead linear, with significant inhibition only observed at concentrations around 2 μM (**Figure 6D, Bottom, Figure S13G**). This deviation in the AD fibrils is notable, as we predicted that our 4R selective TBP-PLP should not be as effective at inhibiting a mixed 3R/4R tauopathy such as AD.

### TBP-PLP Blocks Tau aggregation in human iPSC-derived cortical organoid model

To determine if our PLPs could prevent tau seeding in a model more representative of the human brain, we utilized human forebrain assembloids. These were generated by fusing ventral and dorsal-directed organoids into assembloids, which contain a variety of neuronal subtypes as well as glial cells, including astrocytes ^58–60^.

We treated 6-month-old organoids with sarkosyl-insoluble fractions derived from 9-month-old PS19 mice. These fractions contain highly active tau seeds, which we had previously shown to induce seeding in our planar cell culture biosensor lines. Treatments were administered either with or without 4µM TBP-PLP. For PLP-treated organoids, the PLP was maintained during media changes.

After seven days, the organoids were fixed, sectioned, and stained for MC1, a conformational antibody that specifically detects misfolded tau. In the untreated organoids, strong MC1 signal was observed, particularly in axon-like neuronal processes, predominantly at the outer layer of the organoid (**Figure 7A**).

In contrast, organoids treated with PLP showed a significant reduction in the MC1 staining, indicating that tau misfolding was effectively inhibited in the neuronal processes (**Figure 7B-D**). Notably, the tau detected in this model is likely the endogenously expressed wild-type tau within the neurons.

We trained a machine learning model to identify and partition the signal present in these linear tracts and applied the model to images from multiple sections of four PLP-treated and four untreated biological replicates. The area occupied by this signal was normalized against the perimeter of the organoids, since most of the signal was localized on the periphery of the organoids. An independent t-test on the normally distributed data was significant, with a p-value of 0.017 (**Figure 7E**).

**Figure 7:**
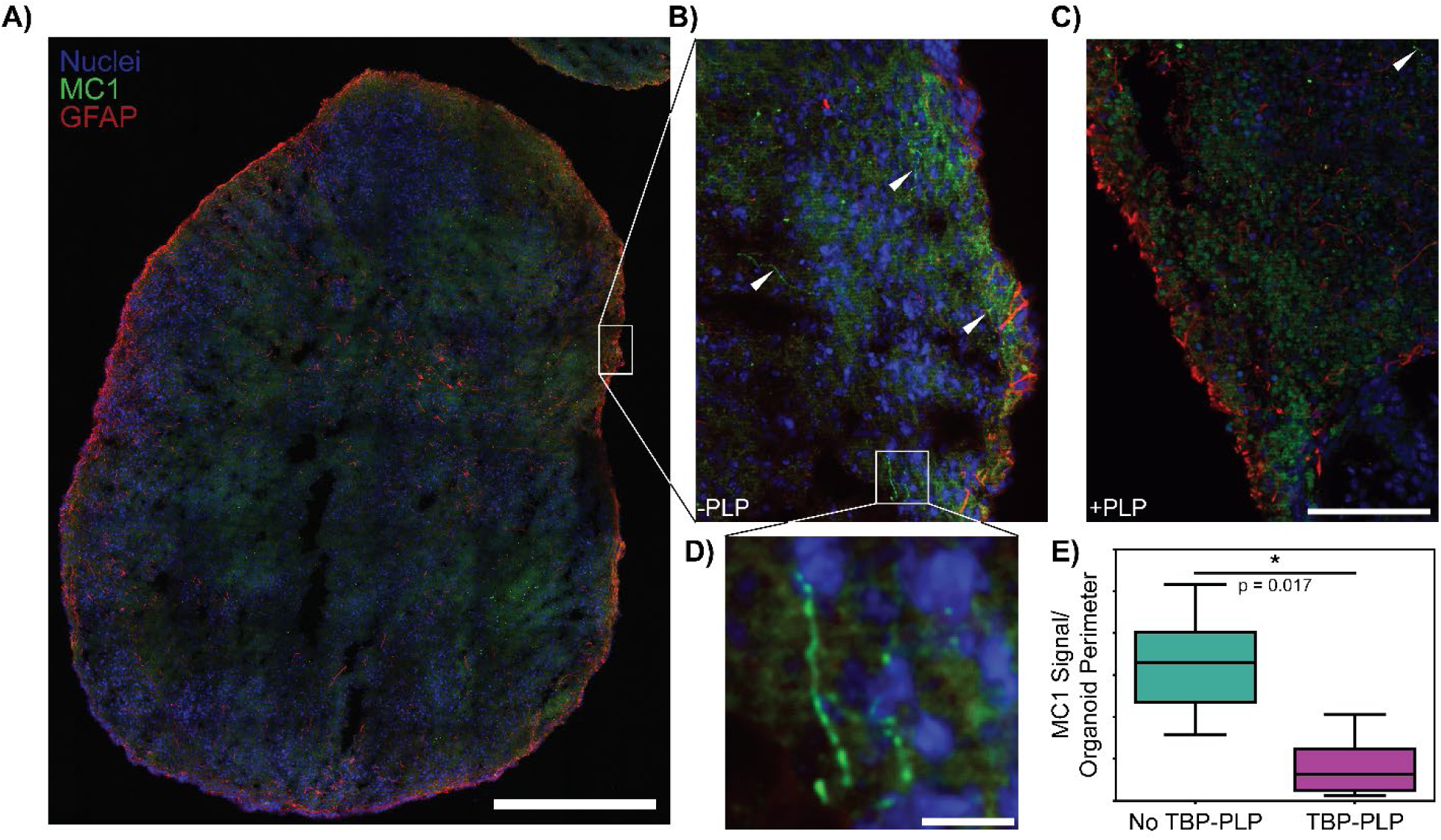
TBP-PLP Prevents Seeding in Human Forebrain Assembloids. Dorsal and ventral assembloids were fused to make forebrain assembloids. A) Assembloids were treated with the sarkosyl insoluble fractions from 9-month-old PS19 mice brains in the absence (A, B, D) or presence (C) of 4 μM TBP-PLP. Organoids were stained with DAPI (Blue), the MC1 tau conformational antibody (green), and GFAP (red). A) Scale Bar: 500 μM. B, C) Scale Bar: 100 μM. D) Scale Bar: 15 μM. E) The area of the signal localized to MC1 positive processes divided by the assembloid’s perimeter showed a significant difference when compared between PLP-treated and untreated (independent t-test, p = 0.017).

## Discussion

Tauopathies, characterized by the accumulation and spread of misfolded tau fibrils, pose a significant health challenge due to their association with neurodegeneration and cognitive decline. The hypothesis that preventing the spread of tau aggregates could halt disease progression is central to our research. To address this, we have developed innovative tools and models to study tau fibril formation, which varies among different tauopathies. Recent advancements in our lab include the resolution of the structure of a minimal tau peptide that selectively seeds 4-repeat (4R) tau isoforms in cellular models^24,25^. We posited that this potent mini-prion could play a different role when incorporated into a multivalent protein-like polymer (PLP) scaffold, effectively occluding fibril ends and preventing further monomer addition after initial binding. This seemingly counterintuitive approach—namely, using a peptide sequence known to fibrillize as an inhibitor—was motivated by our assessment that the PLP is a spherical display of peptides in aqueous solution ^42^, organized as a brush along a flexible polymer backbone, and that this geometry would play a key role in subsequent fibril capping. Indeed, this was borne out in both experimental studies, direct observations and simulations.

PLPs are an attractive drug modality due to their tunable properties for cellular penetration and blood-brain barrier (BBB) permeability ^45,47^. Their multifunctionality also enables their potential as PROTAC molecules. Our findings demonstrate that the 4R-directed TBP-PLP not only inhibits the spread of our model peptide *in vitro* and in cell culture but also effectively blocks the propagation of patient-derived tau fibrils. Importantly, in a human induced pluripotent stem cell (iPSC)-derived brain organoid model, we robustly prevented tau fibril seeding as indicated by the well-characterized MC1 conformational tau antibody. This represents the first application of a bona fide cortical organoid for modeling tau fibril seeding and highlights the potential of this advanced model to further investigate tau pathologies.

Mechanistically, our results show that TBP-PLP directly interacts with tau fibrils both *in vitro* and *in the human brain organoid model*. Transmission electron microscopy (TEM) images are consistent with our assessment that PLPs bind to the ends of fibrils, halting the addition of further tau monomers. Immuno-gold labeling confirms that these aggregates contain PLP, while ThT assays demonstrate that fibril formation is inhibited for at least seven days in vitro. Furthermore, our studies reveal that fluorescently labeled PLP enters cells via direct diffusion and colocalizes with seeded fibrils, preventing the recruitment of additional tau monomers.

A notable challenge in our work was the decreased efficacy of PLP in preventing aggregate formation over extended time points. By maintaining PLP in the media or adding a second dose at 48 hours, we significantly reduced the number of seeded cells to approximately 5%. Additionally, the introduction of a Keap1 PROTAC multifunctional PLP (TBP-Keap1-PLP) demonstrated the potential to prevent tau aggregation even when washed out at six hours post-administration. This area represents a promising avenue for further exploration.

Together, our findings on reduced efficacy at longer time points, the stability of the PLP, and the effectiveness of increased doses or TBP-Keap1-PLP suggest a mechanism whereby TBP-PLP binds to introduced fibrils, preventing the addition of naive monomers. As time progresses, the fibrils are ultimately degraded, but this can lead to an increase in fibril ends. This phenomenon has been previously reported in studies where overexpression of VCP cleared fibrils but resulted in increased seeding due to many smaller fibril fragments^61^. In our system, PROTACs not only facilitated the clearance of these introduced seeds but also allowed for the incorporation of additional TBP-PLP to sponge up the produced fibril ends.

Many therapeutics developed to treat neurodegenerative diseases, particularly those targeting tauopathies, have failed to show significant efficacy in clinical trials. To address these challenges, we developed a novel model system using six-month-old human-induced pluripotent stem cell (iPSC)-derived forebrain assembloids seeded with tau fibrils. Staining with conformational antibodies revealed the presence of aggregated tau in these assembloids. In this more relevant human model, which may offer a better platform for developing drugs with clinical efficacy, we observed that our protein-like polymers effectively blocked MC1 staining, indicating inhibition of tau aggregation. This approach may represent a promising step forward in advancing therapies that translate more successfully from bench to bedside.

Future efforts will focus on reproducing these results in mouse models of tau fibril-induced dementia and enhancing the functionality of these PLPs to improve BBB permeability. By optimizing the design and application of PLPs, we aim to develop effective therapeutic strategies for targeting tauopathies and to potentially halt the progression of these devastating diseases.

## Methods

### Peptide Monomer Synthesis

All peptides were synthesized on Rink resin (0.67mmol/g) using standard Fmoc SPPS procedures on the Liberty Blue Automated Microwave Synthesizer. To generate peptide monomers, amide coupling was performed at the N-terminus, to N-(hexanoic acid)-cis-5-norbornene-exo-dicarboximide (3.0 equiv.) in the presence of HBTU (2.9 equiv.) and DIPEA (6.0 equiv.). The peptide monomers were cleaved from the resin by treating with TFA/H_2_O/TIPS (95:2.5:2.5v/v) for 4h. The crude products were obtained by precipitation in cold diethyl ether, mass confirmed by Bruker Amazon SL and subsequently purified via preparative HPLC.

### Polymer Synthesis

PLPs were prepared via ring-opening metathesis polymerization (ROMP) conducted under nitrogen gas in a glove box. Norbornene conjugated peptide monomers (15.0 equiv., 30mM) were dissolved in degassed DMF with 1M LiCl. Next, the olefin metathesis initiator (IMesH_2_)(C_5_H_5_N)_2_(Cl)_2_Ru=CHPh stock solution (1.0 equiv., 20mg/mL in DMF) was quickly added into the monomer solution. For reaction kinetics, deuterated DMF with LiCl with a hexamethyldisilane standard were used. Here, monomer and polymer olefin peaks were tracked via nuclear magnetic resonance spectroscopy on 400MHz Bruker Avance III HD Nanobay. The solution was left to stir until the full consumption of monomers. Dye tagging was achieved by the addition of 1 eq (30mM) of Cy5.5 linked to norbornene via a six-carbon chain linker with an amide bond. After the polymerization, the polymer solution was purified via preparative HPLC. Finally, the polymer product was obtained by lyophilization.

### Cell Culture and Maintenance

HEK293T and H4 cells were cultured in Dulbecco’s Modified Eagle’s Medium (DMEM) supplemented with 10% Fetal Bovine Serum (FBS) and 1% Penicillin/Streptomycin at 37°C in a 5% CO_2_ environment. Cells were passaged using trypsin equal to 1/10 the volume of the culture vessel and kept under 10 passages to ensure the reproducibility of the data. We have previously described the construction and characterization of the H4 mClover3-Tau187-P301L line used for the majority of the cell-based seeding assays^24^.

### Cell-Based Puncta Seeding Assay

H4 mClover3-Tau187-P301L cells were plated at 20,000 cells/well in a 96-well plate. For all conditions 5 replicates were performed. The following day, varying concentrations of PLP constructs were added to the media for 6 hours unless otherwise noted. After 6 hours, the media was replaced and 2µM jR2R3-P301L fibrils (or disease-derived fibrils at optimized dilutions) were introduced with lipofectamine 2000. 2µM fibril brought to a total volume of 10µL with optiMEM was combined with 1.25µL lipofectamine 2000 in 8.75µL of optiMEM, incubated for 15 minutes at room temperature, added to 1 well. After 24 hours, unless otherwise stated, a minimum of 100 cells were imaged per replicate with confocal microscopy.

### ThT Assay

Thioflavin (ThT) experiments were conducted using a TECAN fluorescent plate reader. Each well of a 384-well plate (Corning, low volume non-binding surface, black with clear flat bottom) contained 25 μM of jR2R3 P301L monomer, 20 μM ThT, 25 μM jR2R3 P301L fibril in a 20mM HEPES buffer (pH 7.4), making a total volume of 30 μL per well. Varying amounts PLP were added before the reaction was initiated.

For degradation assays, 12.5 μM PLP was added to 20mM HEPES pH 7.4, 5 μM Trypsin (from porcine pancreas, Sigma Aldrich, CAS: 9002-07-7) was added and placed into an incubator at 37°C for 30min. 500 μM Inhibitor (Nα-Tosyl-L-lysine chloromethyl ketone hydrochloride, Sigma Aldrich) was then added and allowed to incubate for an additional 30min. 25 μM jR2R3 P301L monomer, 20 μM ThT was added to the mixture. Mixture was then distributed into 5 wells, and 25 μM jR2R3 P301L fibril was added to individual wells for a total volume of 30 μL per well.

The plate reader was set to a temperature of 37°C and allowed time to equilibrate. Subsequently, ThT fluorescence intensity was recorded at excitation and emission wavelengths of 440nm and 485nm, respectively. Measurements were taken every 2 minutes until a plateau in fluorescence intensity was observed. Each experiment was performed with five replicates and repeated three times using independent samples, typically spanning a total time of 24 hours, unless otherwise noted.

### TEM Analysis

For transmission electron microscopy (TEM) analysis, 5 μL of recombinant tau fibril samples were placed onto a glow-discharged copper grid (Electron Microscopy Science, FCF-200-Cu) for 20 seconds before blotting dry with filter paper. The samples were then stained by applying 5 μL of 1.5% (w/v) uranyl acetate solution, immediately blotting dry. An additional 5 μL of uranyl acetate solution was added for 60 seconds, followed by blotting dry. The samples were examined using a Thermo Scientific Talos G2 200X TEM/STEM microscope operating at 200kV and room temperature. The grids were imaged using a Ceta II CMOS 4k x 4k camera at the specified magnifications.

For biotin labeled PLP TEM analysis, the above conditions were used, one without trypsin and one with trypsin. After 24 hours, 10% streptavidin conjugated to gold nanoparticle (Electron Microscopy Sciences, Particle size 6nm) by volume was added and incubated at 4°C overnight. The resulting solution was added to a grid and rinsed 3 times with milliQ water to remove excess gold nanoparticles. The samples were then stained by applying 5 μL of 1.5% (w/v) uranyl acetate solution, immediately blotting dry. An additional 5 μL of uranyl acetate solution was added for 60 seconds, followed by blotting dry.

### Molecular Dynamics

#### Coarse-grained MD simulation on 7P65/PLP complex

We carried out coarse-grained MD simulations using the polarizable MARTINI 2.2P potential.^51,52^ The package GROMACS (version 2023)^53^ was employed. The all-atom structure of the 7P65 protein was coarse-grained using the program martinize2. An elastic network model was applied to preserve the protein structures,^62^ where the neighboring noncovalent CG particles up to a cut-off distance of 0.95nm are structurally constrained using a harmonic bond potential with a force constant of 500 kJ/mol/nm^2^. The MARTINI force field parameters of the polymer backbone have been previously reported by us.^42^ The MARTINI potential has been extensively employed for proteins,^51,52^ polymers,^63,64^ and others.^65–67^ In the MARTINI potential, four non-hydrogen atoms are generally trained as one bead (the 4:1 mapping rule). Four interaction types (polar = P, intermediate polar = N, apolar = C, charged = Q) and some subtypes were introduced to describe the hydrogen bonding capability or the degree of polarity. The MARTINI potential has been proven to be capable of semi-quantitatively reproducing experiments and atomistic simulations beyond the time and length scales that atomistic simulations could afford.^68,69^

The crystal structure of the protein 7P65 was first converted to the MARTINI resolution using the program martinize2. The equilibrated structure of the polymer from the atomistic simulation was converted to the MARTINI resolution using a modified version of the script martinize.py to support the polymer backbone. The protein and the polymer are dissolved in a simulation box with an edge length of 14nm in each dimension. The closest distance between the polymer and 7P65 was around 1.5nm initially (**Figure S17**). The salt concentration was 0.14 M NaCl in addition to the counterions.

The energy of the system was first minimized using the steepest descent algorithm, which was followed by further equilibration of (a) 1ns of simulation using the canonical ensemble (constant number of particles, volume, and temperature, NVT) and the timestep of 10 fs; (b) 1ns of simulation using the isothermal-isobaric ensemble (constant number of particle, pressure, and temperature, NPT) and the timestep of 10 fs; and (c) 15ns of simulation using the NPT ensemble and the timestep of 15 fs. In the production simulations, the recommended parameters^70^ for the MARTINI 2.2 potential were used. Specifically, the short-range Coulomb interactions were calculated up to a cutoff distance of 1.1nm, and the long-range interactions were calculated using the Smooth Particle Mesh Ewald (PME) algorithm.^71,72^ The recommended relativity permittivity of 2.5 was employed for the polarizable water model. The Lennard-Jones (LJ) 12-6 potential interactions were truncated at 1.1nm. The NPT ensemble was applied. The temperatures of water and non-water molecules were separately coupled at 298K using the velocity rescaling method. The isotropic pressure coupling (reference pressure 1 bar, time constant 12.0 ps, compressibility 3×10^−4^ bar^-1^) was employed using the Parrinello-Rahman algorithm. The leapfrog integration time step of 20 fs was employed. The production simulation lasted 20 µs.

#### All-atom MD simulation on the 7P65/PLP complex

Owing to the long-chain feature of the polymers, it is extremely challenging to fully relax the protein/PLP complex using all-atom MD simulations with random initial structures. Inspired by the polarizable MARTINI simulation that the 7P65 protein could form a beta-sheet structure with the peptide sidechains of the polymer (**Figure S7**), we prepared one 7P65/PLP complex with one of the peptide sidechains of the PLP initially aligned with the corresponding 21-amino acid peptide fragment SKDNIKHVPGGGSVQIVYKPV (**Figure S8**). Then, all-atom explicit solvent MD simulation was conducted using GROMACS (version 2023).^53^ Here, the CHARMM 36m potential^17^ was employed, the same as in our previous works on PLPs.^42,45,46,73^ This CHARMM36m potential was specifically optimized to study structureless peptides. The 7P65/PLP complex was dissolved in a simulation box with an edge length of 14nm. A salt concentration of 0.14 M NaCl was employed, along with counterions to neutralize the net charge of 7P65 and the polymer chain.

The system energy was first minimized using the steepest descent algorithm, which was followed by further equilibration using the NPT ensemble of 1ns of simulation with the timestep of 1 fs and 20ns of simulation using the timestep of 2 fs. In these equilibrium simulations, the position restraints were employed on the non-hydrogen atoms of 7P65 and the polymer side chain, which was aligned with 7P65, with the restraint force constant of 1000 kJ/mol/nm^2^. Such restraints were removed in the production simulation below. The three-dimensional periodic boundary conditions were employed. The neighbor searching was done up to 12Å using the Verlet particle-based method and was updated every 20-time steps. The LJ 12-6 interactions were switched off from 10 to 12Å via the potential-switch method in GROMACS. The short-range Coulomb interactions were truncated at the distance of 12Å, and the long-range interactions were calculated using the Smooth Particle Mesh Ewald (PME) algorithm.^71,72^

In the production simulations, the NPT ensemble was employed with the temperatures of water and non-water molecules separately coupled using the Nosé-Hover algorithm (reference temperature 298K, characteristic time 1 ps) and the isotropic Parrinello-Rahman barostat employed with the reference pressure of 1 bar, the characteristic time was 4 ps, and the compressibility of 4.5×10^−5^ bar^-1^. All covalent bonds were constrained, which supported an integration time step of 2 fs. These parameters were recommended for the accurate reproduction of the original CHARMM simulation on lipid membranes,^74^ and have been verified in our simulations on PLPs,^43,45,46,73^ proteins,^69,75–77^ and lipid membranes.^78^

### *In Vitro* Dot Blot Assays

Prior to experiment, all stocks were diluted in 1x Phosphate Buffer Saline (PBS) containing 137mM NaCl, 2.7mM KCl, 10mM phosphate (pH 7.4). 50uL of 10uM jR2R3 P301L Fibril and 2uM R2R3 P301L monomer labeled with FITC fluorophore were incubated with 50uL of 10uM of Biotin labeled JR2R3 P301L species rotating 1 hour at 37C. 50uL of Thermo Scientific™ Pierce™ Streptavidin Magnetic Beads were added to each solution and incubated rotating for 1 hour at room temperature. The beads were washed 3 times with Tris Buffered Saline with 0.1% Tween (TBST) containing 20mM Tris, 150mM NaCl, Tween® 20 detergent: 0.1% (w/v) and eluted with 1mM Glycine pH 2.2. The eluate was subsequently dot-blotted on PVDF membrane and imaged with BIO RAD ChemiDoc MP Imaging System.

Fibrils were extracted from PSP, PS19, CBD, and AD patients’ brain tissue using a sarkosyl treatment and ultracentrifugation. Stocks were diluted with PBS. 20uL of diseased fibrils isolated from human patients were incubated with 20uL of 10uM Biotin labeled JR2R3 P301L species at 37C rotating for 1 hour. 20uL of Thermo Scientific™ Pierce™ Streptavidin Magnetic Beads were added to each solution and incubated rotating for 1 hour at room temperature. The beads were washed 3 times with Tris Buffered Saline with 0.1% Tween (TBST) containing 20mM Tris, 150mM NaCl, Tween® 20 detergent: 0.1% (w/v) and eluted with 20uL of 1mM Glycine pH 2.2. 10uL of eluate was subsequently dot-blotted on PVDF membrane along with unincubated diseased fibril and TBST washes. Tau Monoclonal Antibody (TAU-5; Thermofischer: AHB0042) was incubated at 1 to 1000 dilution in Intercept (PBS) Blocking Buffer with the dot-blotted PVDF membrane overnight shaking at 4 C. Primary incubation was followed by three washes with TBST. Subsequently 680nm anti-mouse secondary antibody was used at 1 to 10,000 dilution was incubated for 1.5 hours and four washes were performed with TBST. The membrane was imaged with BIO RAD ChemiDoc MP Imaging System.

### ICC Experiments

H4 mClover3-Tau187-P301L cells were treated with fluorophore labeled PLPs and after either 6 or 24 hours were fixed with 4% paraformaldehyde for 15 minutes at room temperature. Cells were washed 3x with PBS and placed in blocking buffer (4% BSA, 1% NGS, 0.2% Triton-X100, in PBS pH 7.4) for 1 hour and room temperature. Anti-Rab5 (1:500) or anti Rab7 antibody (1:500) in antibody dilution buffer (2% BSA, 0.5% NGS, 0.2% Triton-X100, in PBS pH 7.4) was added overnight at 4°C. Cells were washed 3x with PBS and secondary antibody (1:500) in antibody dilution buffer was incubated at room temperature for 2 hours. Cells were washed 3x in PBS and imaged.

### ITC Experiments

All isothermal titration calorimetry (ITC) experiments were performed using a Nano ITC (TA Instruments). Prior to the experiments, all buffers were degassed for at least 30 minutes. The experiments were conducted in 1x Phosphate Buffer Saline (PBS) containing 137mM NaCl, 2.7mM KCl, 10mM phosphate (pH 7.4).

Before initiating the experiments, the components were loaded into the Nano ITC and stirred for at least one hour to achieve a stable baseline. A total of 950µL of 500 nM jR2R3 P301L-PLP was added to the gold cylindrical cell of the Nano ITC. Following this, 30 injections of 7.5µL of 5µM jR2R3 P301L monomer were performed, with a 500-second interval between each injection. Identical experiments were performed with jR2R3 P301L fibril samples replacing monomers.

Data analysis was conducted using a combination of NanoAnalyze software (TA Instruments) and custom Python scripts. Control experiments, in which one or both components were replaced with 1X PBS, were performed to determine baseline heats of dilution. These baseline heats were found to be well below those observed in the experimental conditions.

### Preparation of Sarkosyl-Insoluble Fractions

Brain tissue samples were weighed and homogenized in a Dounce homogenizer using 10 volumes of a freshly prepared homogenization buffer containing 0.8M NaCl, 1mM EGTA, 10% sucrose, 0.01M Na_2_H_2_P_2_O_7_, 0.1M NaF, 2mM Na_3_VO_4_, 0.025M β-glycerolphosphate, and 0.01M Tris–HCl at pH 7.4, supplemented with protease inhibitors (Roche, Switzerland). The homogenate was subjected to centrifugation at 20000 rcf for 22 minutes at 4° C, and the supernatant (referred to as SN1) was retained. The pellet was then resuspended in 5 volumes of the same buffer and centrifuged again at 20000 rcf for 22 minutes at 4° C. The second supernatant (SN2) was combined with SN1.

To this pooled supernatant (SN1 + SN2), 0.1% N-lauroyl sarcosinate (sarkosyl; Sigma-Aldrich) was added, and the mixture was agitated on a rotating shaker for one hour at room temperature. The solution was then centrifuged at 125,000 rcf for 63 minutes at 4 °C. After centrifugation, the supernatant was discarded, and the pellet, representing the sarkosyl-insoluble fraction, was rinsed and resuspended in 50mM Tris–HCl (pH 7.4) at a concentration of 200µl per gram of initial tissue. Aliquots of 100µl were stored at −80 °C for subsequent use.

### Human Forebrain Assembloids Generation

Human induced pluripotent stem cells (iPSCs) were donated by Karch ^79^. The iPSC line F12442.4 was cultured in mTeSR Plus medium (STEMCELL Technologies) in 6-well plates (Corning) coated with Matrigel (Corning). Medium was changed daily, and cells were passed with ReleSR (STEMCELL Technologies) when 70-80% confluent.

For the generation of human brain organoids, we followed previously described protocols ^58,59,80^ with some modifications ^60^. On day 0, hiPSC were dissociated into single cells with Accutase (STEMCELL Technologies), then seeded into 96 slit-well plates (S-Bio). 10^4^ cells were seeded per well in mTeSR Plus medium plus 10 nM ROCK inhibitor. After 24h, media was replaced with Neural Induction Medium (NIM), containing Essential 6 (Gibco) as the base, plus 1x Antibiotic-Antimycotic (Gibco), 2.5 uM dorsomorphin (Tocris), 10 uM SB-431542 (Tocris), and 2.5 uM XAV-939 (Tocris). Human cortical organoids (hCO) and human ventral organoids (hVO) were fed daily with NIM from days 1-5, and 5µM IWP-2 was added to medium on days 4-6 only for hVO. On day 6, organoids were transferred to Neural Differentiation Medium (NM), containing Neurobasal-A (Gibco), B-27 supplement without vitamin A (Gibco), 1x GlutaMax, and 1x Antibiotic-Antimycotic, supplemented with 20ng/ml EGF (Shenandoah) and 20ng/ml FGF2 (Shenandoah). hCO organoids were fed daily with NM plus FGF2 and EGF from days 6-15, and every other day from days 16-24. For hVO organoids, NM was also supplemented with 5 uM IWP-2 (Selleckchem) from days 6-11; 5 uM IWP-2, 100 nM SAG (Selleckchem) and 100 nM RA (Sigma) from days 12-14; 5 uM IWP-2, 100 nM SAG, 100 nM RA and 100 nM AlloP (Selleckchem) on day 15; and 5 uM IWP-2, 100 nM SAG, and 100 nM AlloP from days 16-24. Medium was changed daily for hVO from days 6-24. For both hCO and hVO, NM was supplemented with 20ng/ml BDNF (Shenandoah) and 20ng/ml NT3 (Shenandoah) from days 25-43, with medium changes every other day. From day 43 on, hCO and hVO organoids were maintained in NM without additional supplements and fed every 4 days. hCO and hVO were fused at D60 and the fused assembloids used at around D180.

### Immunofluorescence of organoid cryosections

Whole organoids were fixed in PFA 4% for 24h at 4°C, washed 3x with PBS, and then cryopreserved by 48h incubation at 4°C in a solution of 30% sucrose in PBS. Cryopreserved organoids where then embedded in Neg-50 Frozen Section Medium (Epredia), flash frozen in dry ice, and stored at -80° C. Frozen organoids were sectioned in 20µm slices in a *Leica Cryostat* CM1850. For immunofluorescence, fixed organoid cryoslices were incubated in blocking buffer containing PBTA (0.5% BSA and 0.1% Triton X-100 in PBS) plus 5% normal goat serum for 1h at RT. Samples were then incubated with primary antibodies in blocking buffer overnight at 4°C, washed three times with PBTA, then incubated with secondary antibodies in blocking buffer for 1h at RT. Samples were washed three times with PBTA, then mounted with Prolong Diamond Antifade Mountant (Invitrogen) with DAPI for nuclei staining. The following primary antibodies were used: mouse anti-MC1, chicken anti-GFAP (Abcam ab4674, 1:500). The following secondary antibodies were used: goat anti-chicken Alexa Fluor 647 (ThermoFisher A-21449), goat anti-mouse Alexa Fluor 647 (ThermoFisher A-21235). Secondary antibodies were used at a dilution of 1:1,000.

## Acknowledgements

We acknowledge the use of the NRI-MCDB Microscopy Facility and the Resonant Scanning Confocal supported by NSF MRI grant DBI-1625770. N.C.G thanks the National Science Foundation (NSF) for support of work conducted at Northwestern under award number DMR-2403954. Furthermore, M.F. acknowledges support through a NIH predoctoral fellowship. Equipment support was granted through a DURIP/ARO award (for the LCTEM holder) and the National Institutes of Health (S10-OD026871) for TEM camera equipment. This work made use of the EPIC and BioCryo facilities at Northwestern University’s NUANCE Center, supported by the SHyNE Resource (NSF ECCS-2025633), the IIN, and Northwestern’s MRSEC program (NSF DMR-1720139 and NSF DMR-2308691).

## Author Contributions

A.P.L., M.F., J.S.-J., A.D., E.K.R., O.R.S., S.H., B.Q., N.C.G. and K.S.K. designed research; A.P.L., M.F., J.S.-J., A.D., E.K.R., O.R.S, J.Z., J.E.A., M.P.T., E.R.B., E.R.B., C.F., and D.N. performed research; A.P.L., M.F., A.D., J.E.A., and M.P.T. contributed new reagents/analytic tools; A.P.L., M.F., J.S.-J., J.Z., M.T.U., B.F., N.C.G. and K.S.K. analyzed data; and A.P.L., B.Q., N.C.G., and K.S.K. wrote the paper.

