## Supplemental Data for "Protein-Like Polymer for Inhibition of Tau Fibril Propagation in Human-Derived Models of Neurodegeneration"

**Supplementary Information**

### TABLE OF CONTENTS


1. Materials……………………………………………………………………………….3
2. Instrumentation………………………………………………………………………..3
3. Monomer Synthesis and Characterization……………………………………………4
4. Polymer Synthesis and Characterization……………………………………………...7
5. Molecular Dynamics…………………………………………………………………10
6. Binding………………………………………………………………………………...13
7. Residuals………………………………………………………………………….14,16
8. Stability………………………………………………………………………………..15
9. Patient Derived Fibril………………………………………………………………...17

**Supplementary Information**

### Materials and Methods

#### 1. Materials

All materials and reagents, unless otherwise noted, were purchased from commercial sources and used without further purification. *N*-(hexanoic acid)-*cis*-5-norbornene-*exo*-dicarboximide,[1] Cy5.5 norbornene monomer,[2] Biotin termination agent[3] and initiator (IMesH2)(C5H5N)2(Cl)2Ru=CHPh[4] were synthesized according to previous reports.

#### 2. Instrumentation

**2.1 $^1$H Nuclear Magnetic Resonance ($^1$H NMR)**: $^1$H NMR spectra were recorded on a Varian Inova spectrometer (500 MHz) in DMF-$d_7$.

**2.2 Analytical High-Performance Liquid Chromatography (HPLC)**: Analytical HPLC analysis of peptides was performed on a Jupiter 4 Proteo 90Å Phenomenex column (150 x 4.60 mm) using a Hitachi-Elite LaChrom L-2130 pump equipped with UV-Vis detector (Hitachi-Elite LaChrom L2420). The solvent system consists of (A) 0.1% TFA in water and (B) 0.1% TFA in acetonitrile.

**2.3 Preparative HPLC**: A Gilson PLC 2050 purification system was used to purify all peptides. The solvent system consisted of (A) 0.1% TFA in water and (B) 0.1% TFA in acetonitrile.

**2.4 Electrospray Ionization Mass Spectrometry (ESI-MS)**: ESI-MS spectra of peptides were collected using a Bruker Amazon-SL spectrometer configured with an ESI source in both negative and positive ionization mode.

**2.5 Aqueous Phase Gel Permeation Chromatography (Aqueous GPC)**: Aqueous Phase Gel Permeation Chromatography (Aqueous GPC): Aqueous phase GPC measurements were performed with a Tosoh Bioscience TSK-GEL®PWxl-CP column 7.8 mm ID x 30 cm, 10 mm, using 0.05% $NaN_3$ as the mobile phase. Detection consisted of a Wyatt Optilab T-rEX refractive index detector operating at 658 nm and a Wyatt DAWN® HELEOS® II light scattering detector operating at 659 nm.

#### 3. Experimental Methods

##### 3.1 Preparation of Peptide Monomers via Solid-Phase Peptide Synthesis (SPPS) (General procedure)

All peptides were synthesized on CEM ProTide Low Loading resin (0.2 mmol/g) using standard Fmoc SPPS procedures on a CEM Liberty Blue Automated Microwave Synthesizer. Peptide monomers were capped with *N*-(hexanoic acid)-*cis*-5-norbornene-*exo*-dicarboximide using standard SPPS conditions. The peptide monomers were cleaved off the resin by treating the resin with TFA/$H_2O$/TIPS (95:2.5:2.5 v/v) for 2 h. The crude products were obtained by precipitation in cold diethyl ether, and further purified *via* preparative HPLC.

A)

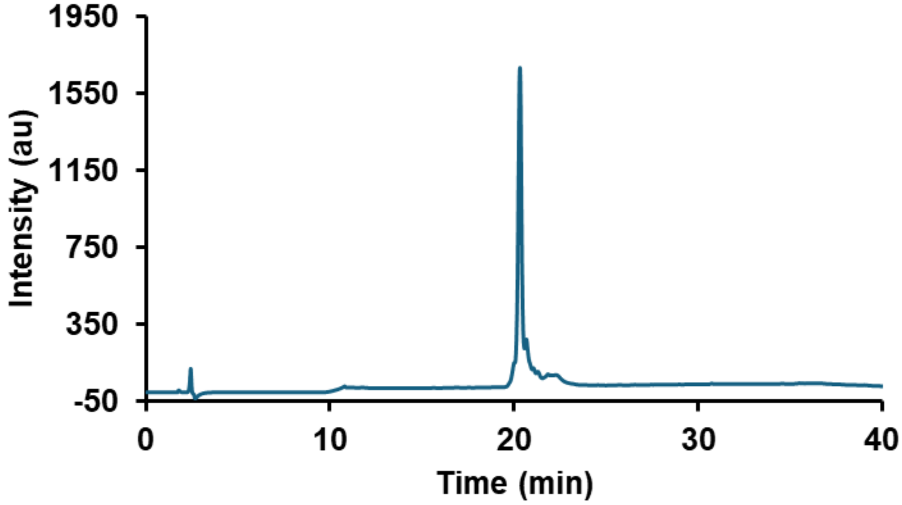
1950
1550
1150
750
350
-50
Intensity (au)
0
10
20
30
40
Time (min)

B)

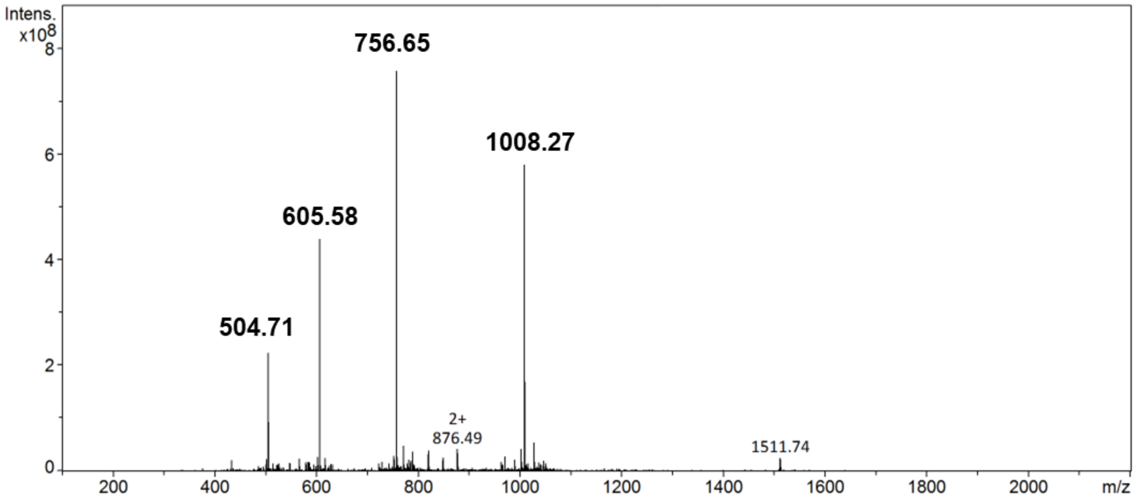
Intens.
x10^8
8
6
4
2
0
756.65
1008.27
605.58
504.71
2+
876.49
1511.74
200
400
600
800
1000
1200
1400
1600
1800
2000
m/z

C)

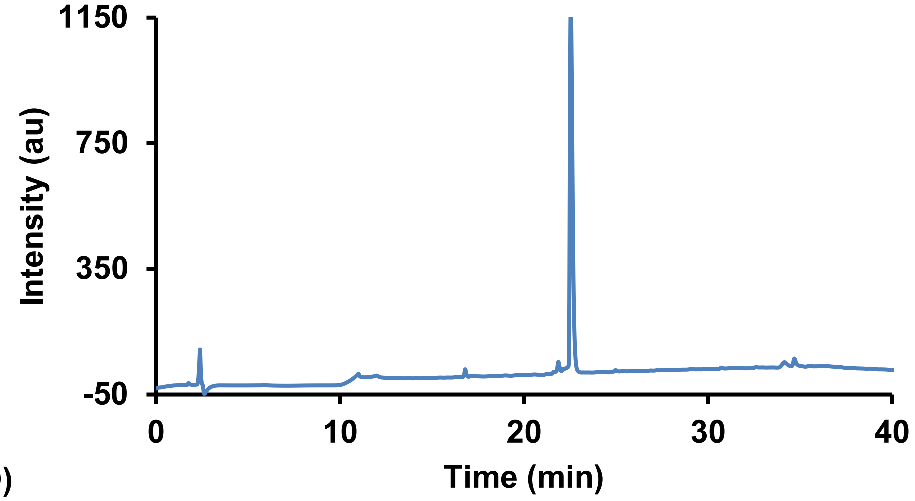


D)

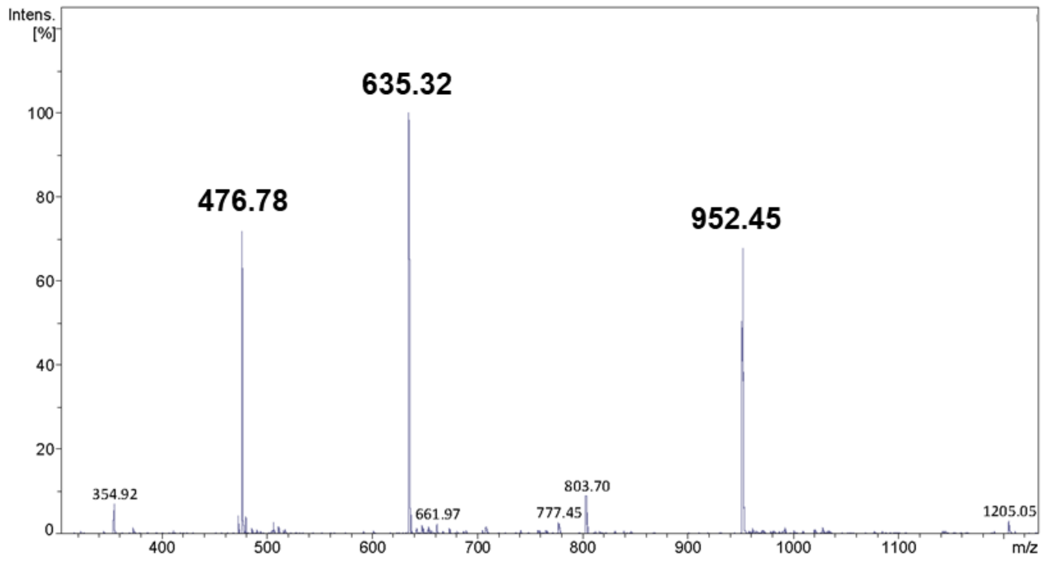


***Figure S1. Characterization of peptide monomers.***
A) Analytical HPLC of jR2R3 P301L monomer NorAha-PGGGSKDINKHVLGGGSVQIVYKPVKK (TBP monomer) after purification. B) ESI-MS of jR2R3 P301L monomer, expected: 3021.57, found: 1008.27 [M+3H], 756.65 [M+4H], 605.58 [M+5H], 504.71 [M+6H]. C) Analytical HPLC of Keap1 monomer NorAha-LDPETGEFLRRRR after purification (Keap1 binder). D) ESI-MS of Keap1 monomer, expected: 1902.96, Found: 980.49 [M+2H], 654.05 [M+3H], 490.81 [M+4H]

##### 3.2 Polymer synthesis and characterization

PLPs were achieved by ring-opening metathesis polymerization (ROMP) under nitrogen gas in a glove box. Norbornene conjugated peptide monomers (20 mg, 15.0 equiv., 30 mM) were dissolved in degassed DMF with 1M LiCl. Next, the olefin metathesis initiator (IMesH2)(C5H5N)2(Cl)2Ru=CHPh stock solution (1.0 equiv., 20 mg/mL in DMF) was quickly added into the monomer solution. The solution was left to stir for 12 h until the full consumption of monomers. In the case of Cy5.5 tagged polymers; this was achieved by the addition of 1 eq (30 mM) of a Cy5.5 linked to norbornene via a six-carbon chain linker with an amide bond. After the polymerization, the polymer solution was precipitated in diethyl ether and was further purified *via* dialysis into deionized water. Finally, the polymer product was obtained by lyophilization.

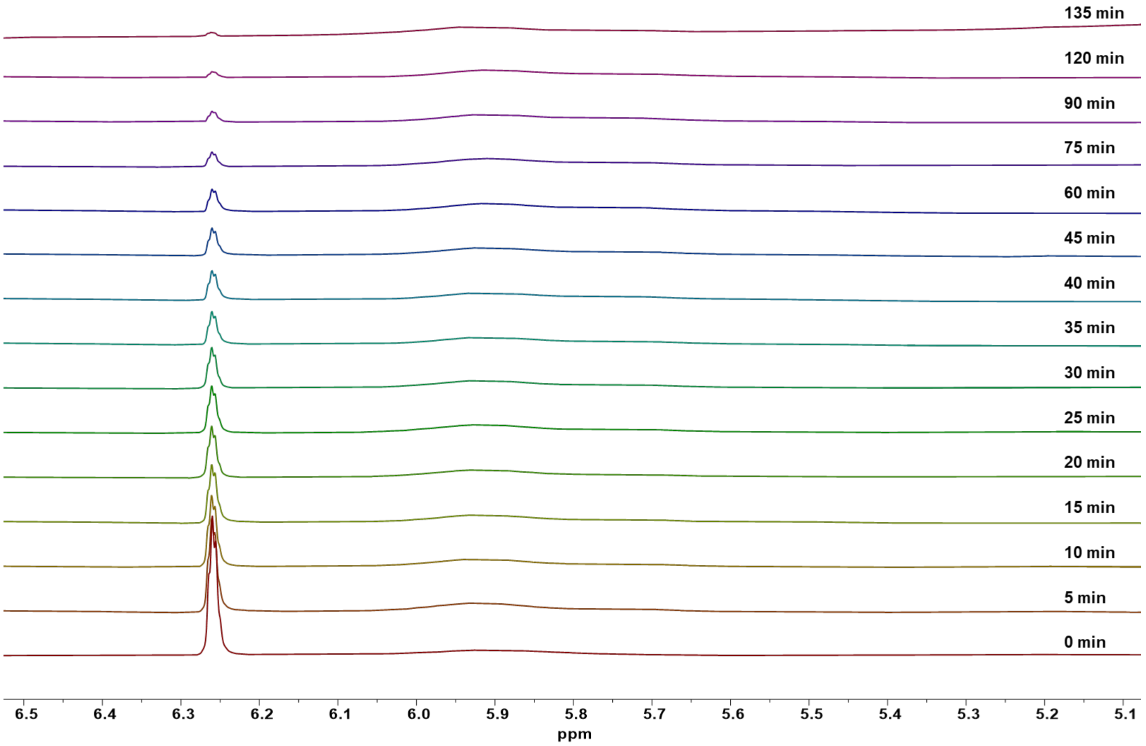


***Figure S2. Polymerization kinetics of jR2R3 P301L peptide monomer.***
Representative polymerization of jR2R3 P301L monomer in 1M LiCl DMF monitored by $^1$H NMR. Disappearance of the norbornene alkene signal at 6.25 ppm was used to determine the progress of the polymerization.

**3.3 Sodium Dodecyl Sulfate Polyacrylamide Gel Electrophoresis (SDS-PAGE)** Protein and polymer samples were denatured and reduced by boiling in Laemmli sample buffer containing SDS. 10 microliters of solutions (unknown for protein, 1 mg/ml for PLP) were loaded onto a precast SDS-PAGE gel and separated by electrophoresis according to molecular weight. Protein bands were visualized by staining with Coomassie Brilliant Blue R-250 and destaining with methanol/water/acetic acid mixture.

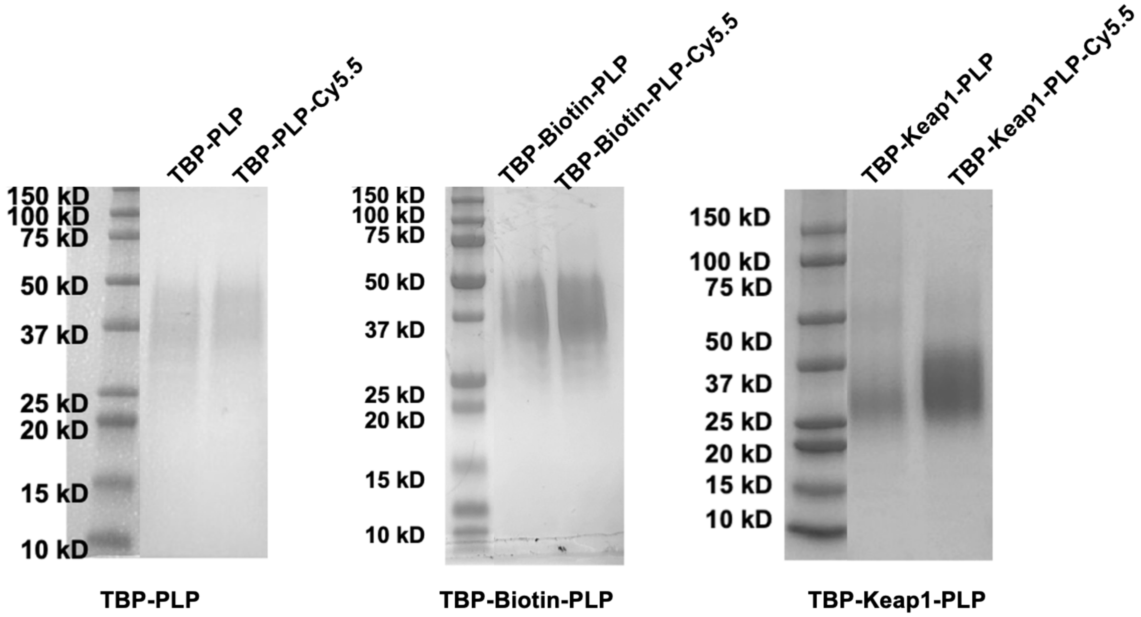


***Figure S3. SDS Page gels used for the characterization of molecular weight of the PLPs used in this study.***
SDS-PAGE of polymers TBP-PLP, TBP-Biotin-PLP, TBP-Keap1-PLP and TBP-Keap1-PLP-Cy5.5.

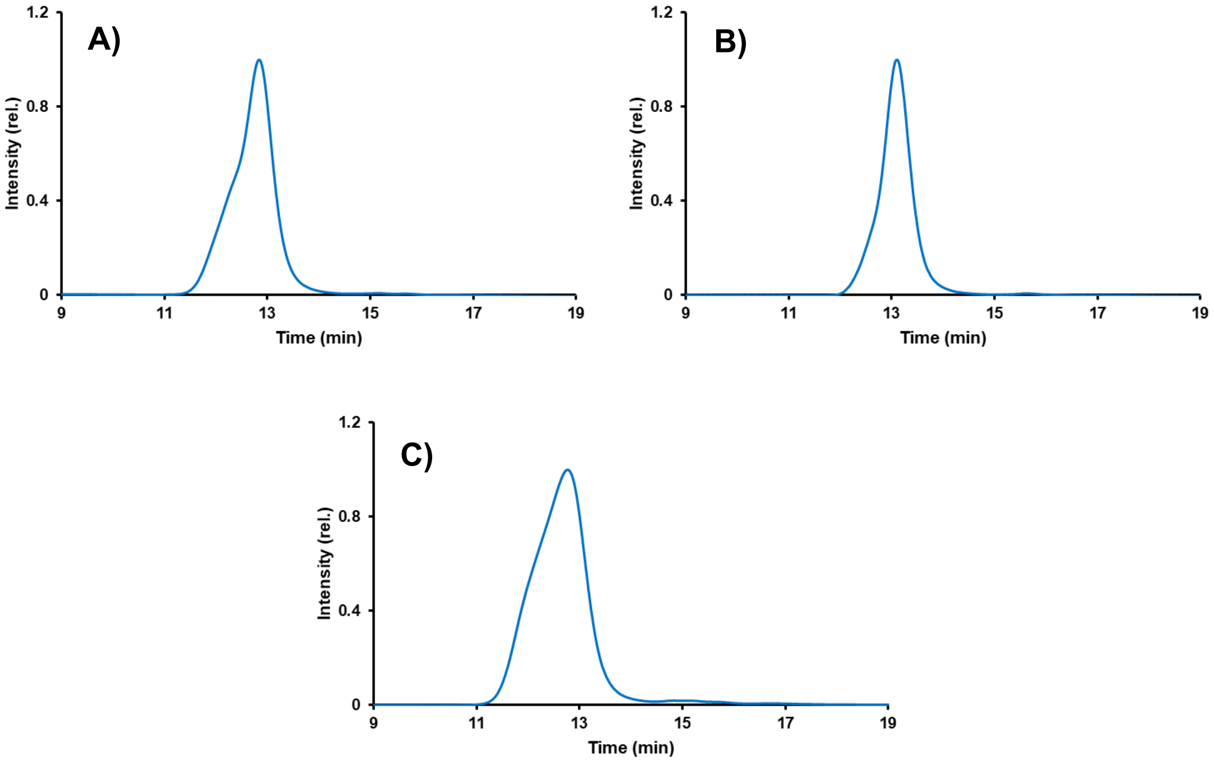


***Figure S4. Molecular Weight determination of PLPs via SEC-MALS***
Aqueous SEC MALS of polymers (A) TBP-PLP, (B) TBP-Biotin-PLP, (C) TBP-Keap1-PLP, run using 0.05 wt% NaN3 in water. Differential refractive index signals are shown.

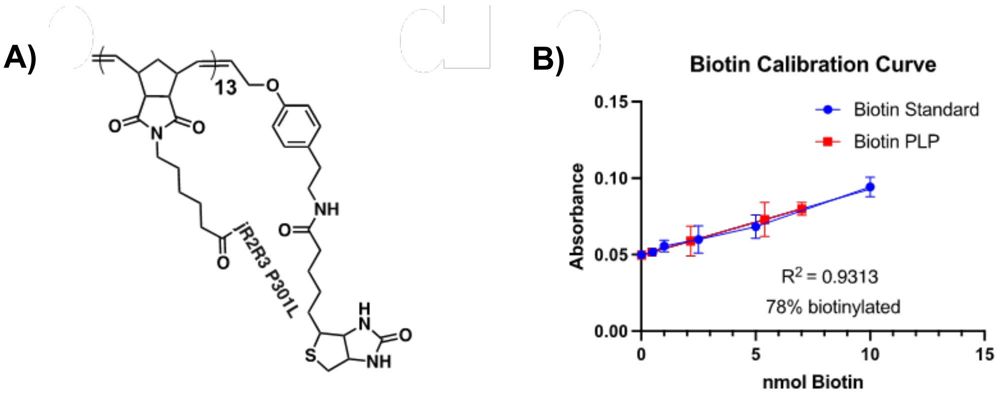


***Figure S5. Characterization of the biotin tagged PLP***
A) Structure of TBP-Biotin-PLP B) Quantification of the degree of biotinylation of the polymer.

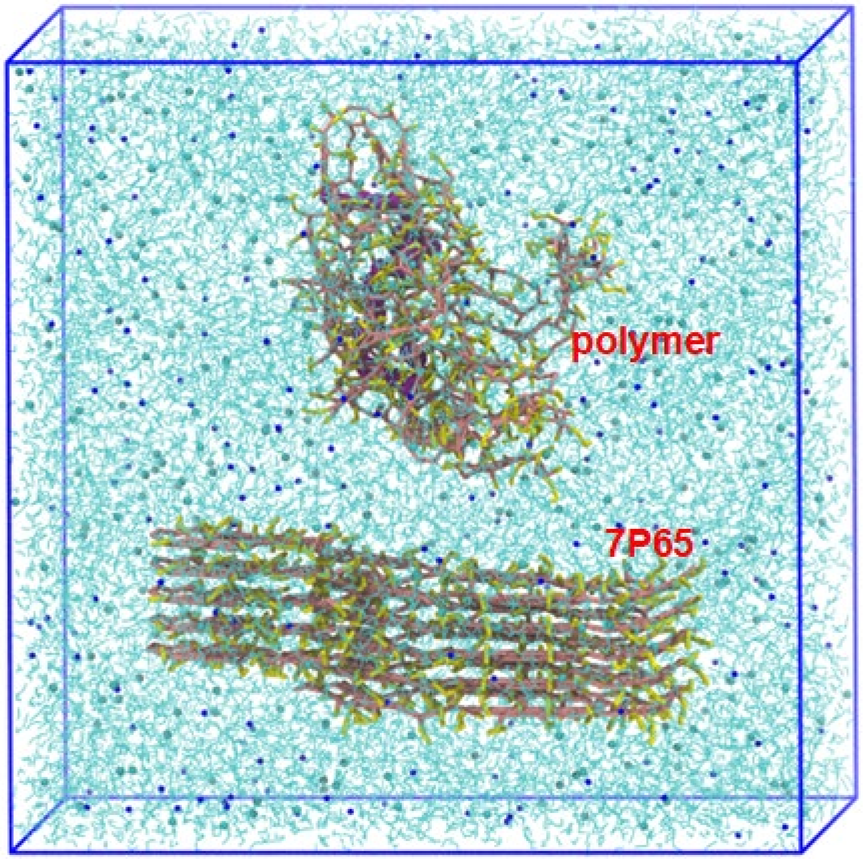


***Figure S6. Starting Structure for Coarse-Grained Simulation***
Initial structure of the MARTINI 2.2P simulation.

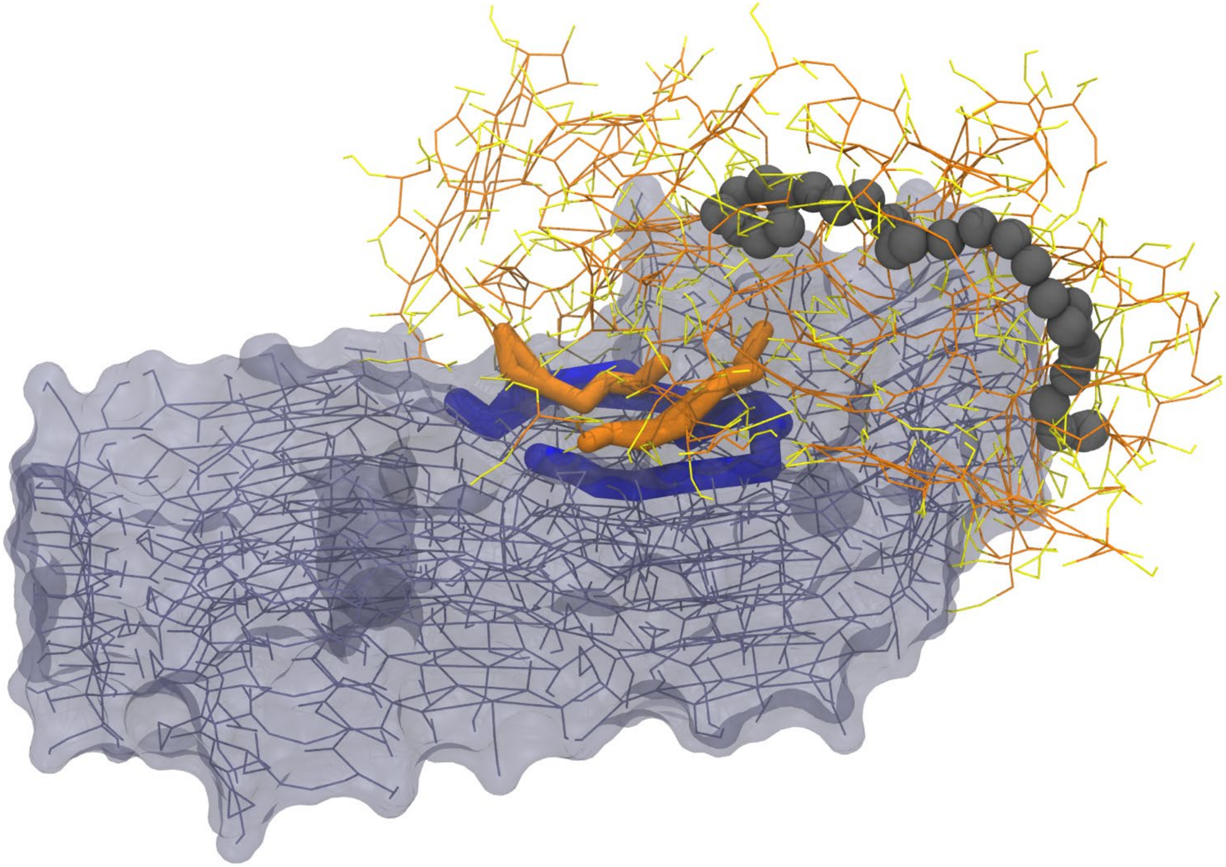

***Figure S7. Coarse-Grained MD of TBP-PLP and PSP Fibrils***
Snapshot from the polarizable MARTINI coarse-grained simulation. The fragment with 21 amino acids SKDNIKHVPGGGSVQIVYKPV is colored with thick blue rods, and the peptide on the PLP sidechain peptide that is aligned with such peptide fragment is highlighted with orange rods. The PLP backbone is highlighted with gray beads. PLP backbone/sidechain beads are colored orange/yellow, respectively.

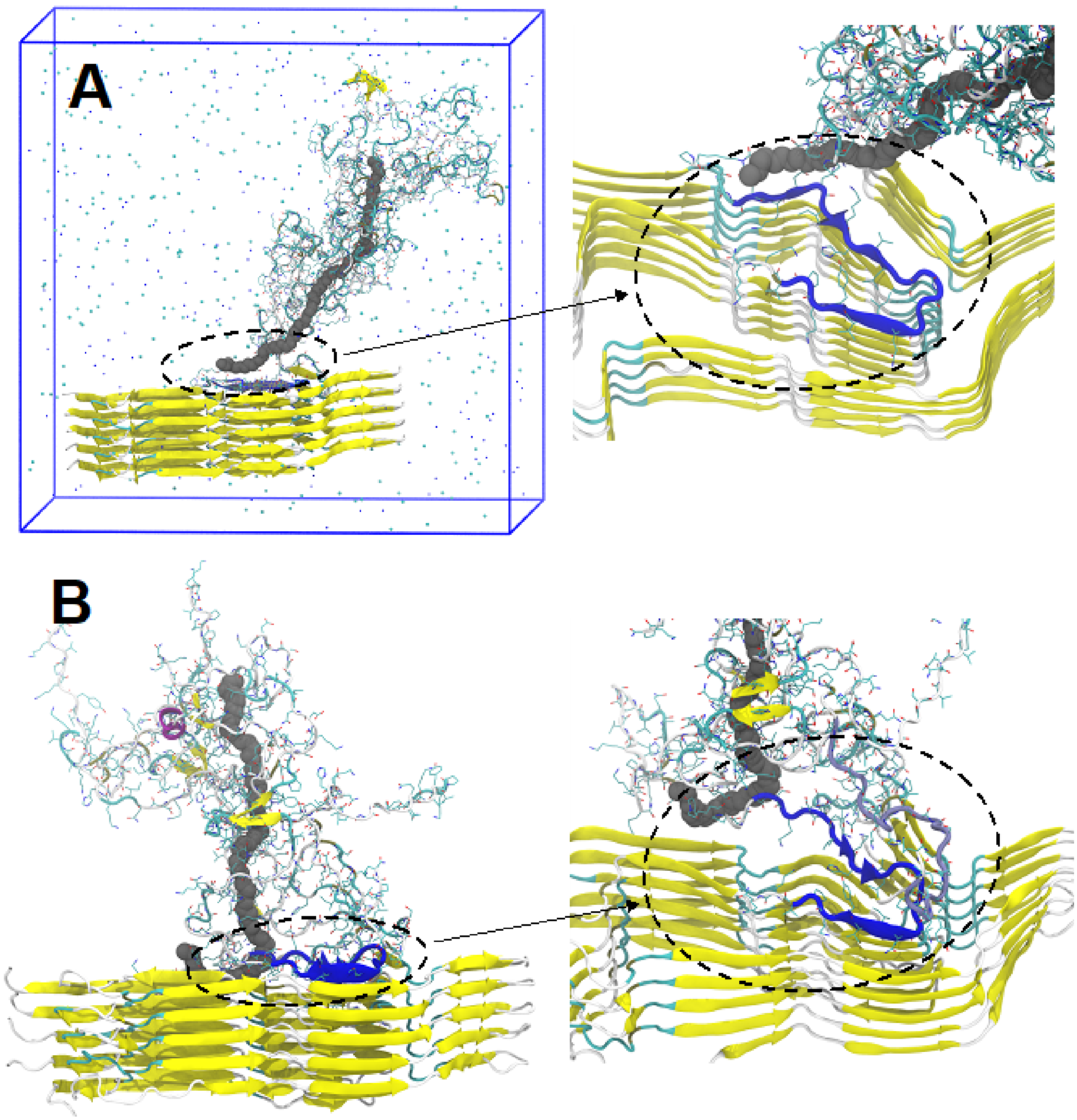


***Figure S8. All-Atom Simulation of the 7P65/TBP-PLP Complex***
(A) Initial and (B) final (time = 300 ns) structures of the all-atom simulation on the 7P65/PLP complex. 7P65 is displayed using the New Cartoon drawing method and colored based on the secondary structures. The 21-amino acid peptide fragment SKDNIKHVPGGGSVQIVYKPV on the PLP that is aligned to 7P65 is highlighted in blue. Demonstrated in the insets are the close views of the 7P65/polymer interface, and in the inset of the final structure, another peptide that is close to the first 21-amino acid peptide fragment is highlighted in light blue. $Na^{+}$/$Cl^{-}$ ions are displayed with blue/cyan dots in the initial structure. Water molecules are omitted from all displays.

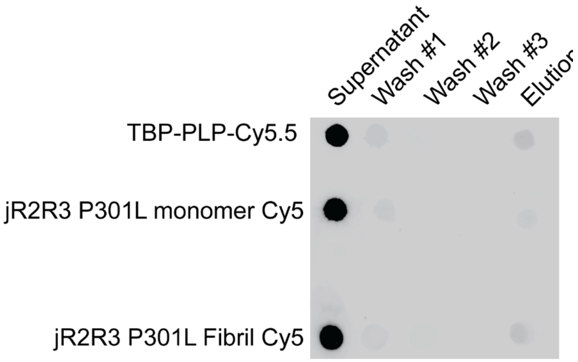


***Figure 9. Control Dot Blots***

TBP-PLP Cy5.5, jR2R3 P301L monomer FITC, or jR2R3 P301L fibril FITC was incubated with streptavidin magnetic beads for 30 minutes and 37˚C and the supernatant, three washes and elution were imaged on a dot blot. No interaction with the beads was observed in these controls.

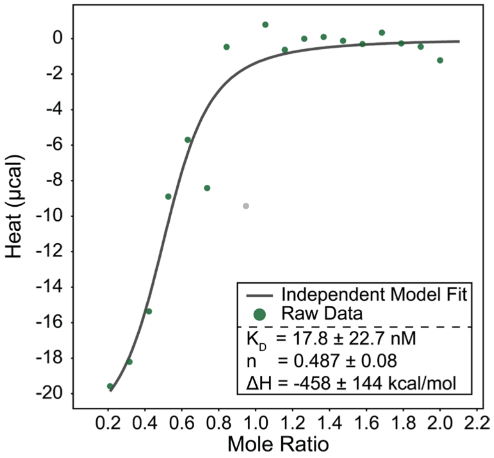


***Figure S10: ITC Measurements of the TBP-PLP/jR2R3 P301L Interaction***

Isothermal calorimetry measurements of the binding between the TBP-PLP (titrate) and jR2R3 P301L monomer (titrant) exhibiting a dissociation constant of 17.8 ± 22.7 nM.

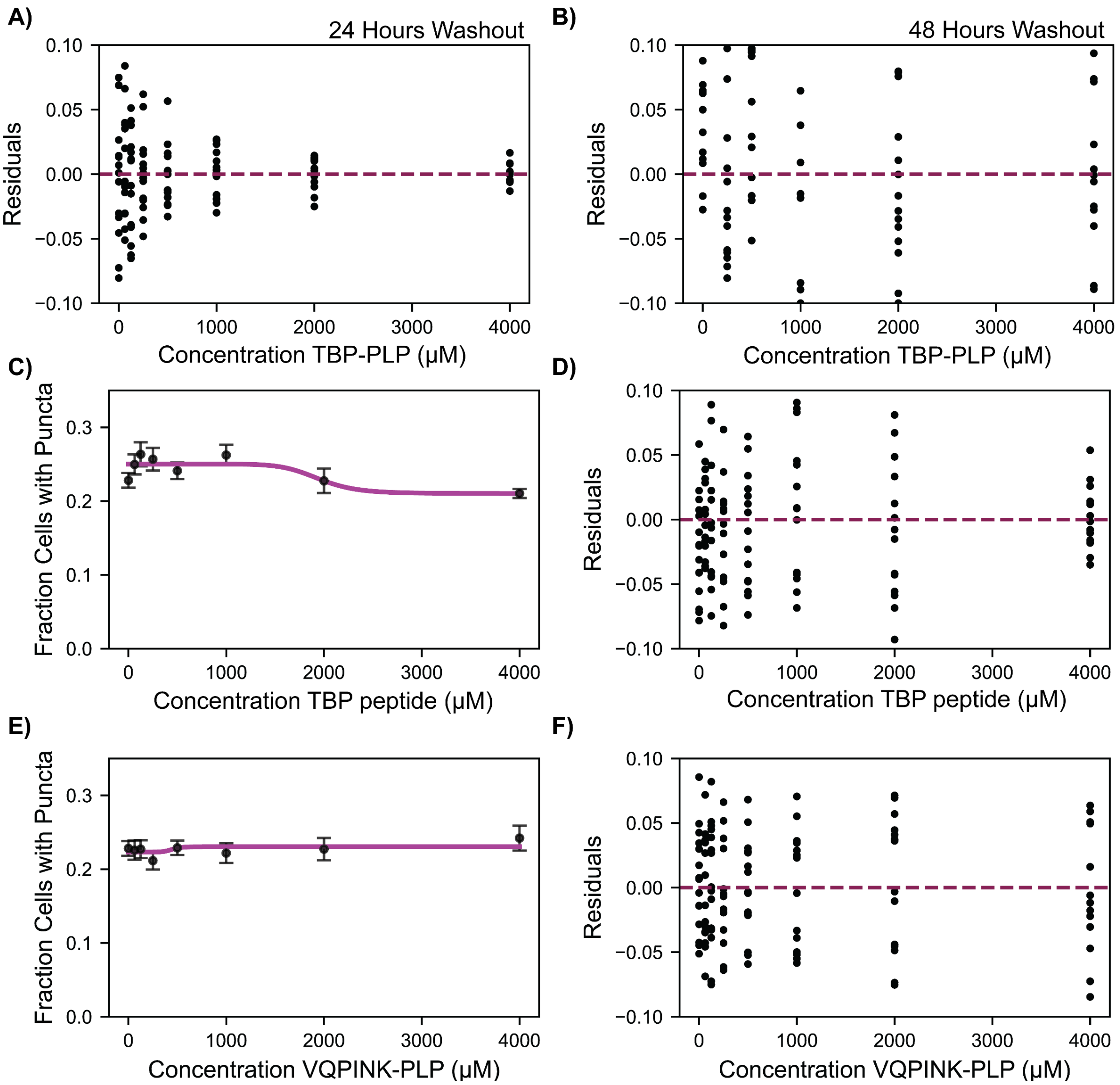


***Figure S11: Residuals, VQPINK-PLP Control and TBP Peptide Controls***
A) Residuals for the 24-hour timepoint in Figure 4C. B) Residuals for the 48 hour timepoint in Figure 4C. C) The fraction of cells containing puncta was plotted against the concentration of TBP peptide (n=15 per condition). Each point represents the average of independent measurements, with error bars showing the standard error of the mean. The data were fit to dose-response curves. D) Residuals for C). E) The fraction of cells containing puncta was plotted against the concentration of VQPINK-PLP (n=15 per condition). Each point represents the average of independent measurements, with error bars showing the standard error of the mean. The data were fit to dose-response curves. F) Residuals for E).

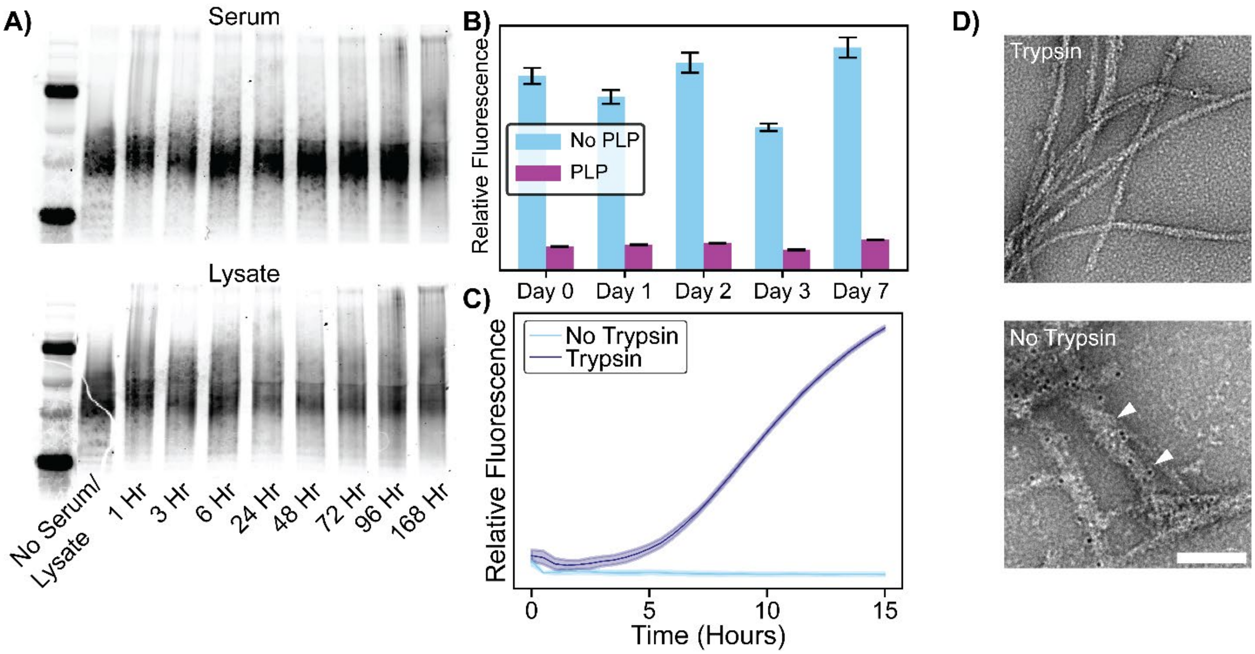


***Figure S12. Stability of TBP-PLP***

A) TBP-PLP was incubated at 37°C with either human serum (top panel) or mouse striatal cell lysate (bottom panel) for 168 hours. Degradation of the PLP was monitored by PAGE at multiple time points. No degradation was observed at any time point in either condition, indicating the high stability of the PLP under physiologically relevant conditions. B) The formation of tau fibrils was monitored using ThT fluorescence over seven days. In the presence of TBP-PLP (purple), no ThT fluorescence was detected, suggesting complete inhibition of tau fibril formation over the entire duration. In contrast, in the absence of TBP-PLP (cyan), fluorescence remained high throughout the experiment, indicating continued fibril formation. C) To assess degradation under forced conditions, TBP-PLP was incubated with trypsin for 60 minutes at 37°C. Trypsin was then inhibited by TLCK, and the treated PLP was introduced to a ThT assay to monitor fibril formation. Trypsin-treated TBP-PLP (purple) showed a significant increase in ThT fluorescence, indicating that degradation of the PLP compromised its ability to inhibit fibril formation. In contrast, untreated TBP-PLP (cyan) continued to prevent fibril formation. D) TEM images from the ThT assays in panel C. In the absence of trypsin treatment, biotin-tagged TBP-PLP was associated with tau fibrils, as indicated by streptavidin immuno-gold labeling (white arrowheads, 100 nm scale bar). However, in trypsin-treated samples, no immuno-gold staining was detected, confirming the loss of PLP binding to fibrils after forced degradation.

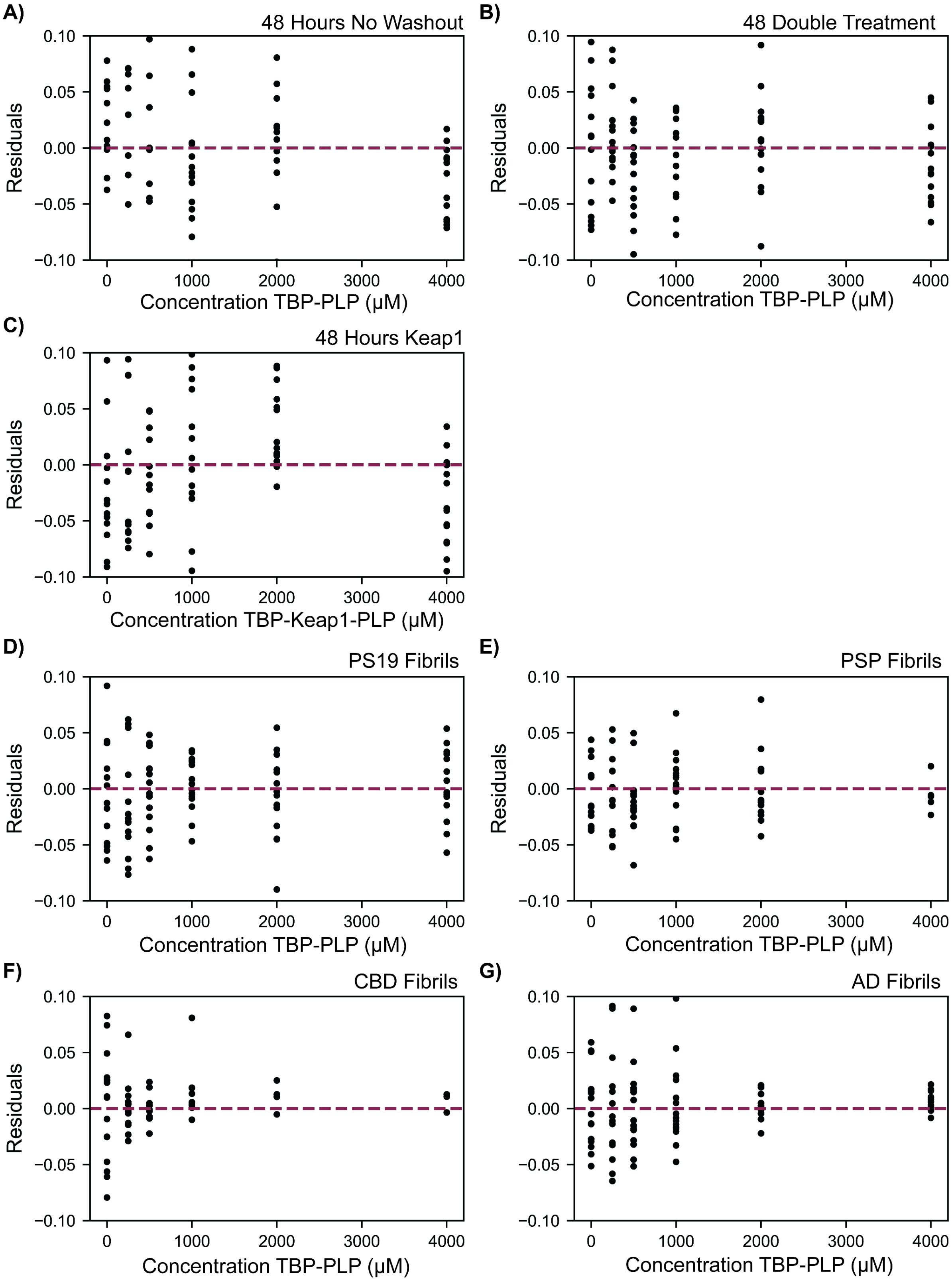


***Figure S13: Residuals of 48 Hour and Patient Fibril Experiments***
A) Residuals of 48 No Washout from Figure 6A. B) Residuals of 48 Double Treatment from

Figure 6A. C) Residuals of 48 hours of TBP-Keap1-PLP treatment from Figure 6B. D) Residuals of PS19 fibrils from Figure 7D. E) Residuals of PSP fibrils from Figure 7D. F) Residuals of CBD fibrils from Figure 7D. G) Residuals of AD fibrils from Figure 7D.

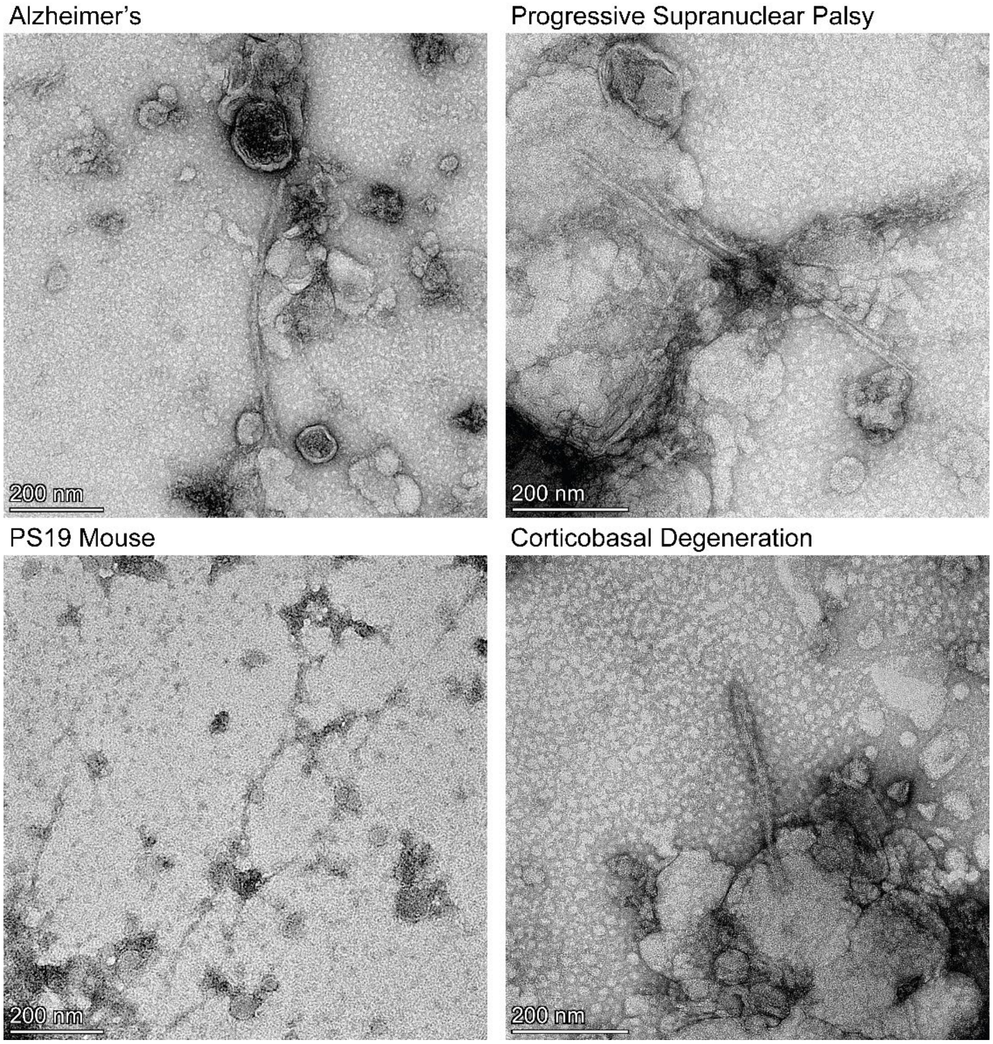


***Figure S14: TEM of Extracted Tau Fibrils***
PSP, PS19, CBD, and AD fibrils were extracted and TEM images were taken to confirm the presence of fibrillar species.

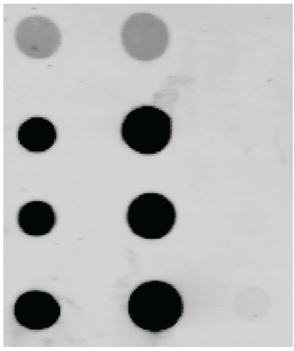


***Figure S15: Control Dot Blot for Disease Fibrils***
PSP, PS19, CBD, and AD fibrils were incubated with magnetic beads in the absence of biotin-tagged jR2R3 P301L-PLP. The first column shows fibrils before incubation with the beads, the second column displays the supernatant containing unbound fibrils, and the last column shows the elution. The blot was stained with a total tau antibody for visualization. Across all conditions, negligible nonspecific binding of disease fibrils to the magnetic-streptavidin beads was observed.
